# Failure to release from the ER translocon induces 60S ribosome degradation

**DOI:** 10.64898/2026.09.26.754687

**Authors:** Zebulon G. Levine, Stuti Khandwala, Christina J. Zhang, Kelsey L. Hickey, Julia Joung, Jingjie Hu, Yi Hua Chen-Neumann, Jingchuan Luo, Anders S. Hansen, J. Wade Harper, Gavin Schlissel, Jonathan S. Weissman

## Abstract

Ribosome malfunction has toxic consequences, but how mammalian cells select ribosomal subunits for degradation remains poorly understood. Here, we develop a pulse-chase CRISPR screening strategy to identify regulators of ribosome turnover and uncover that loss of protein UFMylation promotes ribosome degradation. Normal UFMylation releases the 60S subunit from the endoplasmic reticulum (ER) translocon. Without UFMylation, single-molecule tracking and optogenetic proximity labeling reveal accumulation of ER-associated 60S subunits and their specific degradation. This translocon-associated large subunit degradation (TLSD) is independent of the lysosome and, in our human cell model, is only active in nondividing, contact-inhibited cells. Degradation requires the poorly characterized protein R3HDM4, which we term TLSD1. Upon UFMylation loss, TLSD1 redistributes to the ER and promotes 60S degradation. These findings identify a cell-state-dependent pathway for clearing ER-retained 60S subunits. We propose translocon release acts as a checkpoint for ribosome function, explaining TLSD1’s genetic links to neuronal health and erythropoiesis.

## Introduction

Ribosomes are macromolecular complexes that catalyze protein translation, the most energetically expensive and error-prone step of gene expression.^1,2^ Effective translation requires the ribosome to find the correct reading frame and coordinate nascent chain synthesis with chaperones and targeting machinery; failures both reduce functional protein output and can generate toxic misfolded proteins.^3–7^ To prevent this, eukaryotic cells carefully monitor both ribosome biogenesis and translation through quality-control pathways.^6,8^ During assembly, nascent ribosomes of both the large (60S) and small (40S) subunits must pass through multiple quality-control checkpoints;^8–10^ particles that fail are targeted for degradation to keep the translating pool of ribosomes functional. Translation is also surveilled by quality control pathways.^6,15,16^ Slow decoding or stalled ribosomes trigger translation termination, ribosome splitting, and degradation of both the nascent chain and mRNA.^6,7,15^

Because ribosomes are costly to build, they are typically recycled in translation failures rather than degraded; however, a defective ribosome could seed a cascade of further translation failures. Mature ribosomal subunits are long-lived, with mammalian ribosome half lives ranging from several days to more than a week,^11–14^ potentially allowing damage to accumulate.^17–20^ Consistent with this possibility, a recent study found that ribosome collisions increase with a ribosome’s molecular age,^21^ and translation dysfunction is known to increase with organismal age and appear early in neurodegeneration.^22–25^ The stoichiometry of ribosomal proteins and oxidation of rRNA are also altered with age and disease, raising the possibility that defects in quality control of ribosomes themselves contribute to translation dysfunction.^22,25–27^ However, the machinery that monitors mature ribosome integrity and selects ribosomes for degradation remains poorly understood.

The mechanisms responsible for ribosome degradation are far less studied than those governing biogenesis, especially in mammals.^28^ Ribosomes are degraded through both autophagic and proteasome-dependent pathways. Ribosomes traffic to the lysosome upon nutrient starvation via autophagy of ribosomes (ribophagy), which has also been observed upon translation and oxidative stress;^29–33^ however, it remains controversial whether this reflects targeted degradation, because ribosomes often undergo nonselective engulfment, or bystander autophagy, during degradation of other cargo.^29,30^ Non-lysosomal ribosome degradation is also known, including the late-acting cytosolic pathways that degrade 40S and 60S subunits bearing nonfunctional rRNAs, termed nonfunctional rRNA decay (NRD).^34–42^ These pathways, discovered and characterized in yeast,^34,37,39–41^ act at late cytoplasmic stages of assembly, and only the 40S degradation pathway has been demonstrated to recognize fully matured ribosomes. Recently, this 40S arm of NRD has been characterized in human cells,^35^ where homologs of the yeast pathway similarly ubiquitinate nonfunctional or initiation-stalled 40S subunits; notably, the pathway appears to have diverged in humans, with metazoan-specific protein RIOK3 acting as the ubiquitin-recognizing effector.^35,43–46^ Mature 60S degradation, by contrast, remains enigmatic in mammals, and the large subunit NRD pathway genes lack clear human homologs.

Ribosome degradation is likely to be of particular importance in tissues with long-lived, slowly dividing cells. In rapidly dividing cells, biogenesis replenishes the ribosome pool with quality-control during assembly, and cell division dilutes older ribosomes.^29,47,48^ In nondividing cells, biogenesis is balanced with degradation to maintain ribosome level. Without dilution through division, defective ribosomes have the potential to accumulate if they are not degraded. Because studies have largely used rapidly dividing models, ribosome homeostasis in nondividing mammalian tissues remains underexplored, as does whether quiescent or post-mitotic cells safeguard their translation machinery differently from cells that dilute damage through division.

Here, we systematically mapped genes contributing to ribosome turnover in a quiescent human cell model. We combined genome-wide CRISPR interference (CRISPRi) screening with pulse-chase labeling of endogenously HaloTagged ribosomal proteins to identify genes that, upon knockdown, modulate the ratio of new to old ribosomes for each subunit. We then developed a complementary screening strategy to distinguish effects on subunit synthesis from effects on degradation. Our screens recovered known ribosome turnover and biogenesis pathways and found that both ribosomal subunits are degraded faster in response to loss of UFMylation, a ubiquitin-like modifier pathway that acts on post-termination large subunits at the endoplasmic reticulum (ER) translocon. Focusing on 60S subunits, we combined single-molecule ribosome tracking and optogenetic proximity labeling, we elucidated a previously-unrecognized ribosome degradation pathway, which we term translocon-associated large subunit degradation (TLSD). TLSD degrades 60S ribosomes in nondividing cells and requires the poorly characterized protein R3HDM4, which we propose renaming TLSD1. We propose that release from the ER translocon serves as a checkpoint for large subunit function, with failed release promoting degradation of dysfunctional subunits.

## Results

### Systematic identification of genes involved in ribosome homeostasis

To systematically screen for pathways regulating ribosome turnover in nondividing cells, we turned to hTERT-RPE1 (RPE1) cells, a telomerase-immortalized retinal pigment epithelium cell line that maintains a near-euploid karyotype with intact quality control pathways including p53 signaling.^49–51^ These cells can be induced to quiescence by contact inhibition, reduced serum and growth factor signaling, or a combination^52,53^ (Figure 1A)-indeed, combining these two signals provided <3% positivity in an EdU staining assay for entry through the cell cycle^54^ in a 24-hour period, while approximately two-thirds of RPE1 cells stained positive for DNA synthesis when grown under standard conditions. We used CRISPR-Cas9-mediated homology-directed repair to endogenously tag RPL29 and RPS3 with HaloTag7 (HaloTag) (Figure 1B, Supplementary Figure S1A), allowing us to use these “RiboHalo” cell lines to implement a previously-reported pulse chase assay to track ribosome turnover in individual cells.^29,55^ These ribosomal proteins were chosen both because they have previously been validated to incorporate well into the ribosome when HaloTag-modified,^29,56^ and because they do not show evidence of exchanging once incorporated, turning over at a rate consistent with the bulk of the ribosome.^11,57^ We saw no defects in expression of these tagged proteins relative to untagged copies by immunoblot (Supplementary Figure S1A). The HaloTag protein covalently and irreversibly binds cell permeable small molecule dyes,^58^ allowing fluorescent labeling of all tagged ribosomal proteins in the cell at that timepoint-washing allows removal of unincorporated dye, so newly synthesized proteins remain unlabeled (Figure 1C). At the end of the experiment, a second dye can be added, labeling new proteins. This allows single cell measurements of new ribosomal proteins, old ribosomal proteins, and the ratio between them.^29,53,55^

**Figure 1:**
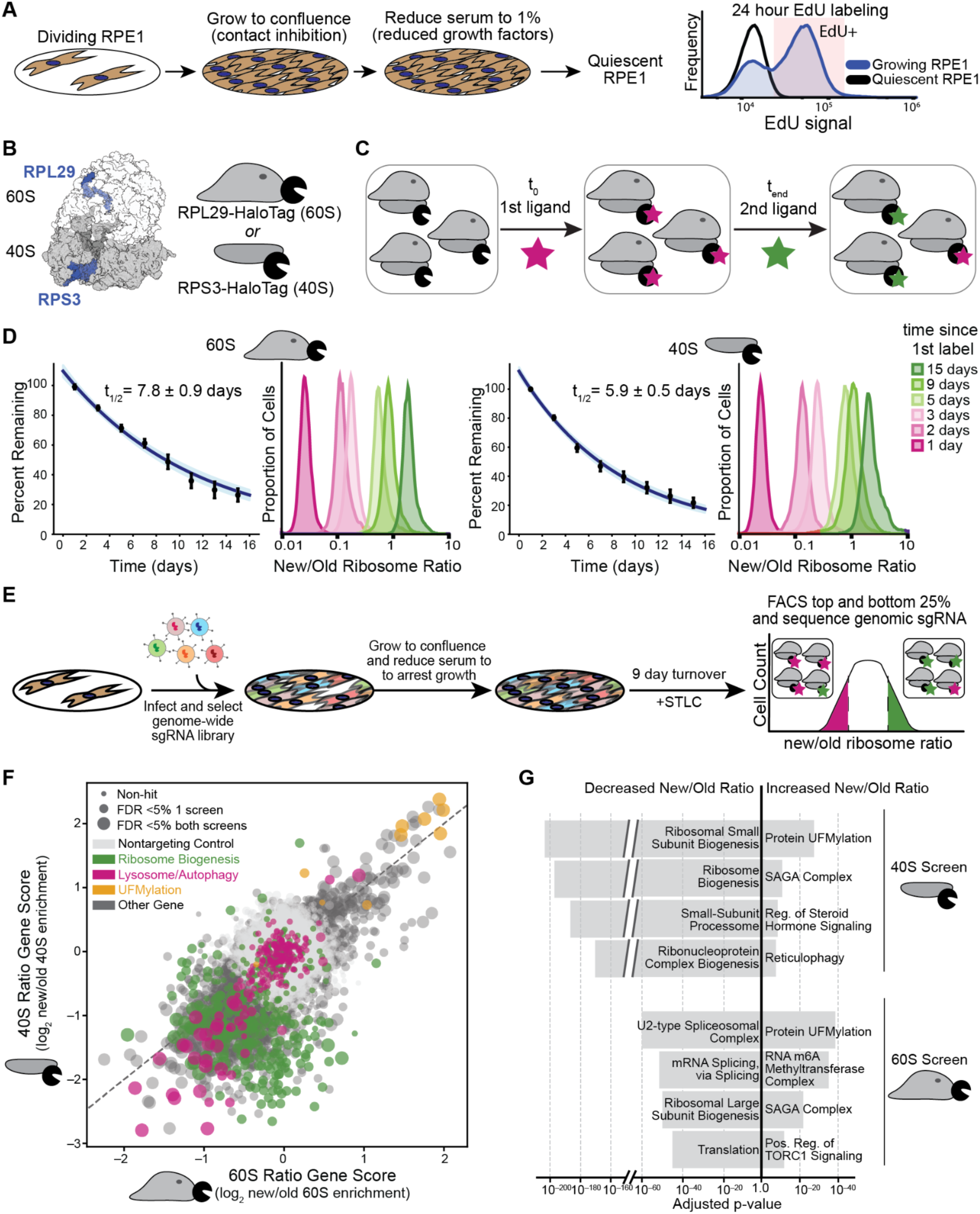
Systematic screening uncovers regulators of ribosome synthesis and degradation in nondividing human cells A) RPE1s enter quiescence from a combination of contact inhibition and serum reduction. Right: flow cytometry after 24 hours incubation with EdU to stain for active DNA synthesis under quiescent or growing conditions. B) HaloTag7 was added to the endogenous locus of RPS3 (40S subunit) and RPL29 (60S subunit) in separate clonally isolated RPE1 cell lines. C) RiboHalo turnover assay schematic. JF549-Halo (magenta) is added at the start of the assay, labeling pre-existing ribosomes, then dye is removed. JF646-Halo (green) is added at the end to label newly synthesized ribosomes. D) Turnover curves and per-cell new/old ribosome ratios, split by large subunit (left) and small subunit (right). Histograms colored by day of acquisition after initial labeling. Turnover curves show mean and standard error of 3 replicates each timepoint, normalized to singly-labeled controls; blue lines and t_1/2_ are exponential decay fit with 95% confidence interval. E) Schematic of genome-wide screen on new/old ribosome ratio. STLC is a mitotic kinesin inhibitor that will trigger M-phase arrest and prevent division of quiescence escapees. F) Scatter plot showing gene knockdown effect on new/old ribosome ratio for small subunit (y-axis) and large subunit (x-axis). Positive scores indicate increased new/old ratio upon gene knockdown. Point size represents whether gene passes >5% false discovery rate threshold. Genes colored by pathway. G) GO categories enriched for increasing or decreasing new/old ribosome ratio upon knockdown. Adjusted p-value is Benjamini-Hochberg corrected.

To validate our single-cell assay for ribosome turnover, we performed labeling experiments using flow cytometry. As the majority of ribosomal proteins are incorporated into ribosomes and short-lived outside of the ribosome,^18,59,60^ we reasoned that this strategy would provide an accurate approximation for ribosome turnover rate. We used Halo-alkane-modified versions of Janelia Fluor 549 (JF549-Halo) and Janelia Fluor 646 (JF646-Halo) as our “old” and “new” dyes as these provide a high signal to noise.^61^ Fitting a single-exponential decay function to this data yielded half-lives of 7.8±0.9 days for the large subunit and 5.9±0.5 days for the small subunit (Figure 1D), which are similar to rates obtained from nondividing mouse tissues via mass spectrometry with heavy amino acids,^11,12^ arguing the RPE1 system provides a reasonable model for nondividing mammalian tissues. Importantly, for each time point, comparing the ratio of new to old ribosome signal provided a narrow distribution that was dependent on time since labeling (Figure 1D).

This new/old ratio provides an internally normalized metric we used to perform a genomewide CRISPRi screen (Figure 1E).^62–64^ We isolated RiboHalo reporter clones expressing Zim3-dCas9 CRISPRi machinery and introduced a 3 guide-per-gene genomewide CRISPRi guide library (Supplementary Table 1) into dividing RPE1 RiboHalo large or small subunit screening cell lines. After infection and selection, we grew the pooled screen to quiescence and carried out a 9-day turnover experiment in duplicate for each subunit. During the turnover experiment, we added S-trityl-L-cysteine (STLC) as an inhibitor of mitotic kinesin KIF11; this arrests M-phase cells and leads to cell death, preventing overgrowth of quiescence “escapees”.^53,65^ In parallel, ribosomes were labeled for turnover in conditions which allowed RPE1 growth and collected after 3 days to compare effects on ribosome turnover rates in dividing versus quiescent cells. For both conditions, fluorescence-activated cell sorting (FACS) was used to purify cell populations in the top and bottom 25% of new/old ratio. Unsorted cells from mid-selection and the end of the experiment were collected to measure gene knockdown effects on cell growth and survival. The sequencing-based log_2_ fold change in relative guide RNA abundance after amplification from genomic DNA was used to obtain gene-level effects of knockdown on ribosome turnover (new/old ratio score) and growth, with nontargeting control guides used to measure statistical significance (Tables S1–S2, Figure 1F, Supplementary Figure 1C–E).^66,67^

Comparing the screening outcomes from the 3-day dividing cell turnover experiment to those from quiescent cells demonstrated the necessity of carrying out this assay in a nondividing model. During sorting, we noted that the new/old ribosome ratio histogram was left-skewed in growing cells (Supplementary Figure 1B). We reasoned that cells with low new/old ribosome ratio likely had reduced growth rate and thus did not dilute the old ribosomes during division. Consistent with this, the new/old ratio scores from the growing cell screen were better correlated with growth phenotype (Pearson r of 0.57–0.70) than with new/old ratio scores from the quiescent cell state (Pearson r of 0.28–0.40) (Supplementary Figure 1C). Growth effects were highly correlated (Pearson r of 0.80) between reporter cell lines and essential genes depleted in both, consistent with the tagged ribosomal proteins not changing cell state (Supplementary Figure 1D). In contrast to these growth effects and the new/old ratio scores from growing cells, new/old ratio scores from quiescent cells have lower correlation between subunits, suggesting distinct degradation and synthesis patterns for each subunit. Additionally, essential genes have both positive and negative scores (Supplementary Figure 1E). Taken together, this data suggests that the new/old ribosome ratio reports on a distinct process from cell division in quiescent cells.

Focusing on the quiescent screens, we examined genes expected to have effects on ribosome synthesis or degradation (Figure 1F, Supplementary Figure 1F). We find negative new/old ratio gene scores (low new/old ratio) for ribosome biogenesis gene knockdown, which lowers the level of new ribosomes, as well as autophagy and lysosome gene knockdown, which increases old ribosome levels if cells are engaging in ribophagy. Prior research suggests that “bystander ribophagy” during nutrient deprivation is a major source of ribosome turnover.^28–32^ We next explored if the low serum conditions used in our studies led to higher autophagic flux. We used a global autophagy reporter and compared change in flux upon loss of core autophagy genes FIP200 and ATG7 under different growth conditions (Supplementary Figure 1G).^68^ Adjusting from growing cells to quiescence increased autophagy by similar amounts to overnight amino acid starvation, confirming that our screening system had high basal autophagy consistent with the strong ribophagy signal.

Three intriguing trends emerged from our genome-wide data. The first was the observation that while knockdown of biogenesis genes for either subunit led to reduced new/old ratio in the 40S screen, the large subunit only strongly responded to genes altering 60S biogenesis directly, with limited effects of 40S biogenesis knockdown (Figure 1F, Supplementary Figure 1F). Second, while literature reports both proteasomal and lysosomal ribosome degradation pathways,^28–44^ in the quiescent screen, we see limited effects of proteasome knockdown relative to lysosome gene knockdown (Supplementary Figure 1F). Last, knockdown of genes involved in the transfer of ubiquitin fold modifier 1 (UFM1) increased the new/old ribosome ratio for both subunits. Protein UFMylation scored as the most enriched pathway for increasing new/old ribosome ratio, while ribosome biogenesis dominated effects for decreasing new/old ribosome ratio (Figure 1G, Table S3). We found this intriguing: while UFMylation is known to directly modify the ribosomal large subunit,^69,70^ it is not known to play a role in ribosome level control, although it has been implicated in autophagy of the ER (ERphagy)^71^ and promoting the degradation of nascent chains derived from ribosome stalling and collisions at the ER translocon.^72–75^ However, the above screen could not determine if UFMylation’s effect on ribosome levels was due to biogenesis or degradation.

### Systematic classification of ribosome degradation or biogenesis genes

Altering ribosome biogenesis or degradation rates can have the same effect on new/old ratio by changing either the numerator or the denominator; however, they have opposite effects on total levels (Figure 2A). Indeed, simulating changes in degradation and synthesis rates suggests that comparing new/old ratio to absolute abundance should clearly distinguish effects on synthesis from effects on degradation (Supplementary Figure 2A). In order to classify candidate genes based on their specific impact on biogenesis versus degradation, we cloned a sublibrary of the 715 top-scoring genes from the genomewide screen (Supplementary Figure 2B). Each reporter cell line was infected with this sublibrary and sorted after a 9-day turnover experiment based on three different metrics, collecting top and bottom 25% for each: total new ribosome level, total old ribosome levels, and the new/old ribosome ratio used in the genome-wide screen (Supplementary Tables 4–5).

**Figure 2:**
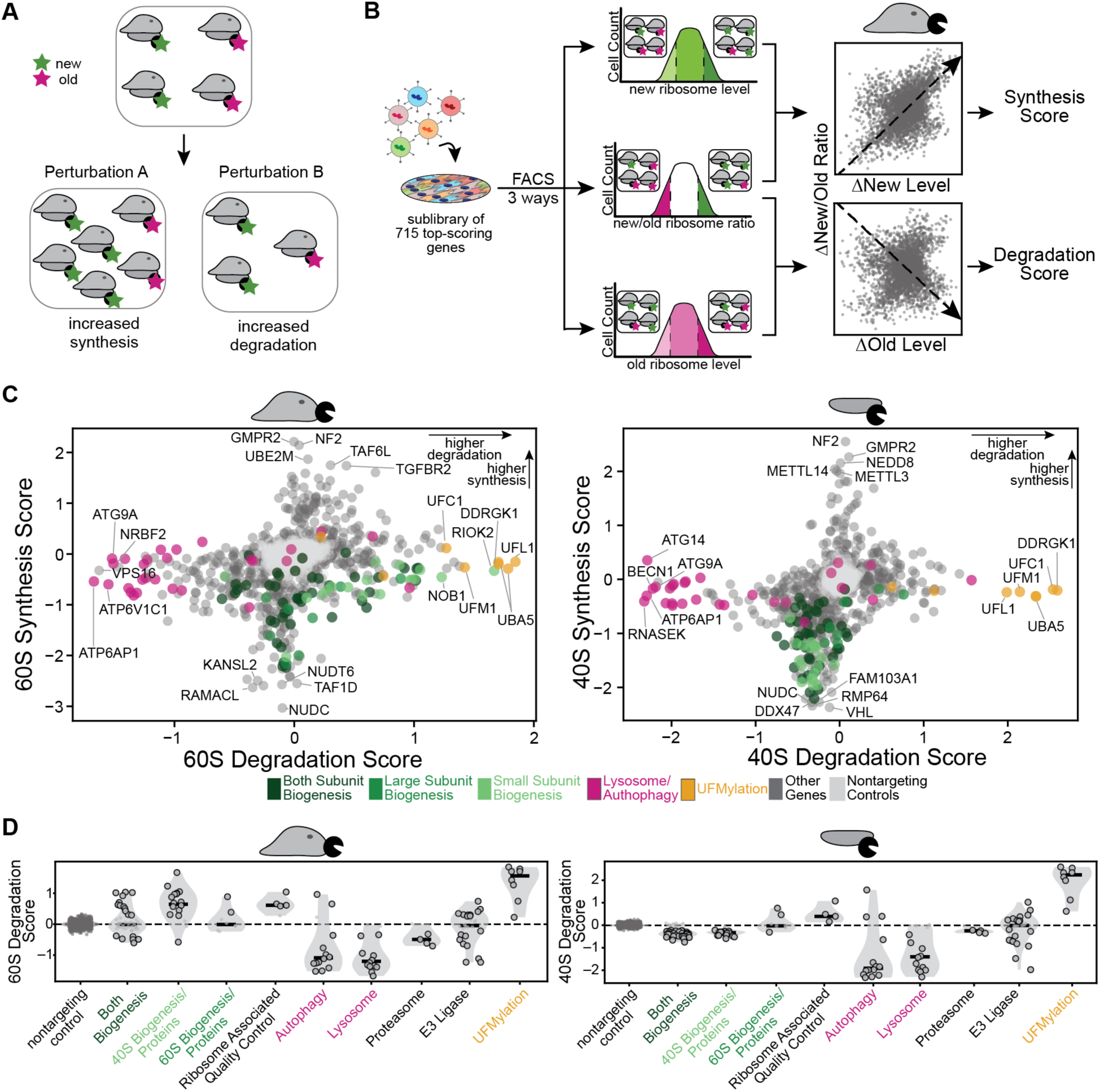
Score-based classification of ribosome degradation and synthesis shows accelerated ribosome degradation upon loss of UFMylation. A) Perturbing biogenesis (X) or degradation (Y) can have the same effect on new to old ribosome ratio, but would have different effects on ribosome biogenesis B) Schematic of screening strategy and score development. Screen carried out as in Figure 1E, but sorting on old or new ribosome level in addition to new/old ratio. Synthesis and degradation scores based on distance along lines shown in sgRNA-level data plots. C) Gene-level scores for large subunit (left) and small subunit (right). Scores represent weighted log_2_ fold change (see Methods) in synthesis (y-axis) or degradation (x-axis); positive scores indicate increased synthesis or degradation respectively upon knockdown. D) Effects of gene categories on degradation score for large subunit (left) and small subunit (right). Circled dots pass 5% false discovery rate compared to nontargeting controls by Mann-Whitney U-test.

When we compared the log_2_ fold sgRNA enrichment between high and low new/old ribosome ratio to enrichments based upon changes in new ribosome level or old ribosome level, we observed the data formed an X-shaped pattern (Figure 2B, Supplementary Figure 2C), as we would predict from genes primarily altering either degradation or synthesis. Ribosome biogenesis knockdown causes new/old ribosome ratio and ribosome abundance to change in the same direction, while knockdown of ribosome degradation causes discordant changes in ratio versus abundance. Consistent with our simulation, we saw higher dynamic range along the “degradation axis” when comparing old ribosome levels relative to new/old ratio, and higher dynamic range for synthesis when looking at new ribosome levels (Supplementary Figure 2D). We captured these effects as sgRNA and gene level “scores” for ribosome synthesis and degradation (Supplementary Figure 2C, Supplementary Tables 4–5).

Autophagy and ribosome biogenesis cleanly separate between degradation score (autophagy) and synthesis score (biogenesis), having negative scores in their respective category and providing confidence in our metric (Figure 2C–D, Supplementary Figure 2E). Consistent with the pro-growth role of ribosome biogenesis,^76^ growth effects correlated highly with synthesis scores but not degradation scores (Supplementary Figure 2F). Similar to the genomewide effects, we noted that small subunit-specific biogenesis genes have limited synthesis score magnitude for the large subunit, but knockdown in biogenesis of either subunit decreased 40S synthesis scores (Figure 2C–D, Supplementary Figure 3A–B). This is consistent with findings in yeast showing asymmetric responses to perturbation of proteins of each subunit,^77,78^ and recent findings in human cells showing small subunit degradation in response to impaired large subunit biogenesis.^44^

Intriguingly, while large subunit synthesis scores showed limited response to small subunit biogenesis knockdown, a subset of 40S biogenesis genes induced large subunit degradation upon knockdown (Figure 2C–D, Supplementary Figures 2E, 3A–B). We noted that these 40S biogenesis genes (RIOK2, NOB1, RIOK1, TSR1, PNO1) were all late cytoplasmic 40S maturation factors,^8,79,80^ and hypothesized that late cytoplasmic small subunit synthesis failure may promote mature 60S subunit degradation. While similar effects have been observed in yeast,^10,77^ this feedback was not thought to occur in mammals.^79,80^

### Loss of large subunit UFMylation leads to degradation of both ribosomal subunits

Strikingly, loss of UFMylation emerged as the most prominent pathway perturbation that selectively accelerated ribosome degradation, with strong effects on both the large and small subunits and little effect on biogenesis (Figure 2C–D, Supplementary Figures 2E, 3A). We confirmed that UFMylation loss led to degradation of the whole ribosome rather than just our HaloTag reporters using stable isotope labeling using amino acids (SILAC) with mass spectrometry. We mimicked our HaloTag labeling approach, fully labeling cells with heavy arginine and lysine during growth to quiescence, then shifting to light amino acids and collecting at 3 and 8 days in samples with either control guides (sgNT) or knockdown of DDRGK1 (sgDDRGK1), a component of the E3 transferase complex for UFM1 (Figure 3A–C, Supplementary Figure 4A, Table S7).^81,82^ As predicted from the chase experiment setup, old (heavy) protein levels decreased over time, while new (light) protein levels increased (Supplementary Figure 4B, Table S8). The change in levels of light protein level between the two timepoints provided a metric of synthesis rate, while the change in old protein level provided a metric of degradation rate. We could therefore directly compare changes in synthesis and degradation between genotypes. Comparing both levels and rates of synthesis and degradation globally, we noted correlation between the total protein level changes and synthesis rate changes, while degradation rate changes, as expected, clustered separately, with anticorrelation to changes in new protein levels (Supplementary Figure 4C, Table S9). Degradation rate increases upon UFMylation loss for both ribosomal subunits is highly specific and is not seen across organelles or other complexes (Figure 3B and Supplementary Figure 4D–E). Notably, the proteomic analysis reveals stabilization of the core Sec61 translocon (Supplementary Figure 4F), suggesting that ribosomes are degraded separately from the translocon.

**Figure 3:**
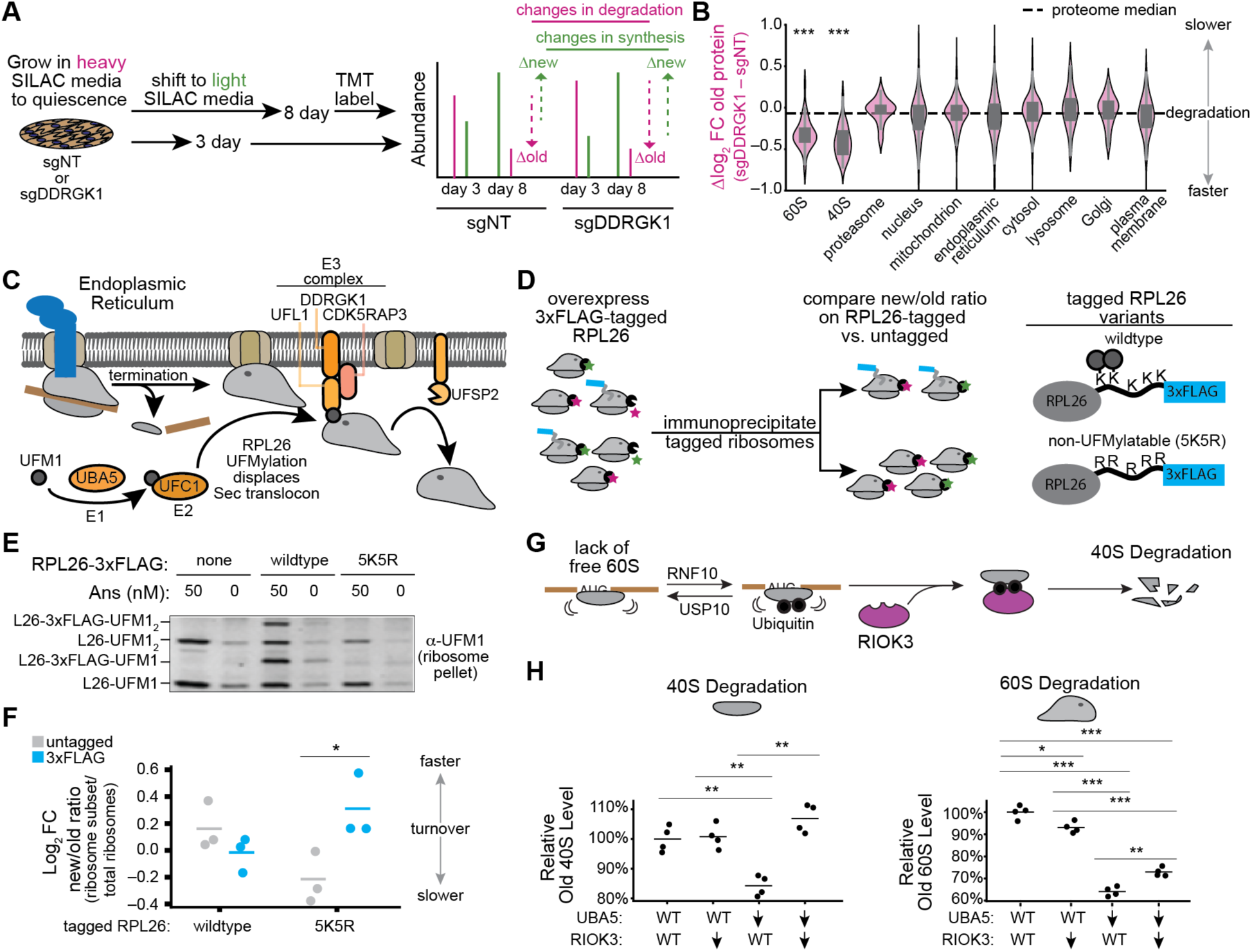
Failure to UFMylate RPL26 leads to degradation of both 40S and 60S ribosomal subunits. A) Schematic of SILAC-tandem mass tag (TMT) pulse labeling experiment for sgNT (control) and sgDDRGK1 (UFMylation knockdown) cells. B) Difference in log_2_ fold change old proteins over time between genotypes for categories of proteins (listed in Table S8). ***: p <0.001, Mann-Whitney U versus rest of proteome. C) UFMylation of the ribosome at the ER relies on E1-E2-E3 cascade to modify RPL26, displacing the large subunit from the translocon. RPL26-UFM1 is held at the ER until deUFMylation by UFSP2. D) Schematic of non-UFMylatable ribosome pulldown assay. 3xFLAG-tagged RPL26 is overexpressed in cells with RPL29-HaloTag and endogenous untagged RPL26. RiboHalo pulse-chase labeling allows comparison of RPL26 variants that can or can’t be UFMylated. E) UFM1 immunoblot with ribosome collision induction using low-dose (50 nM) anisomycin for 15 minutes before collection. RPL26-3xFLAG tagged constructs expressed in cells as indicated. F) Quantification of new/old 60S ribosome ratio at 5 days after initial labeling (relative to signal from ribosome pellet before pulldown). Horizontal line is mean, circles are individual data points. n=3 per condition. *: p <0.05, Welch’s t-test. G) Schematic of RNF10-RIOK3 pathway for small subunit degradation in response to lack of free 60S subunits availability for translation initiation. H) Change in relative old ribosome level (left: 40S, right: 60S, 5 days after initial labeling) upon CRISPRi knockdown of RIOK3, UBA5, or both, measured using flow cytometry (points show median, normalized to average value for no knockdown). *: p<0.05, **: p< 0.01, ***: p< 0.001, Welch’s t-test.

The primary target of the UFMylation pathway is the C-terminal tail of RPL26 (uL24) in the ribosomal large subunit, which is modified at the ER with the small ubiquitin like protein UFM1.^69,70,83^ The UFMylation pathway uses an E1-E2-E3 cascade to modify the large subunit on the ER translocon after either normal termination at a stop codon or 60S-40S splitting in response to ribosome stalls and collision;^75,84–86^ subsequent conformational changes cause the E3 ligase to bind the transferred UFM1 and displaces the large subunit from the Sec61 translocon (Figure 3C).^84,85^ Without this transfer, biochemical evidence shows that the 60S-translocon complex is stable and 60S release is inefficient.^84,85,87^ To test if RPL26 UFMylation, as opposed to UFMylation of other targets, is responsible for accelerated large subunit degradation, we generated 3x-FLAG tagged variants of RPL26 with either wildtype sequence or a non UFMylable mutant we termed 5K5R, which replaced five C-terminal lysines including the sites of UFMylation with the non-nucleophilic charge-preserving arginine (Figure 3D).^69^ We overexpressed these variants in RPL29-HaloTag cell lines, so that both normal, untagged ribosomes and our tagged RPL26 variants would coexist in every cell. Inducing ribosome collisions with low-dose translation elongation inhibitor anisomycin, which is known to promote RPL26-UFMylation,^74,75^ caused the appearance of bands corresponding to mono-and di-UFMylated RPL26 (Figure 3E). Expressing FLAG-tagged wildtype RPL26, but not the 5K5R mutant, caused the appearance of higher-molecular-weight UFM1 bands, consistent with modification of the wildtype tagged RPL26 but not the 5K5R mutant. We did a five day pulse-chase labeling of these ribosomes using RPL29-HaloTag to track turnover under quiescent conditions and separated the FLAG-tagged and untagged ribosomes after ribosome isolation via immunoprecipitation (Supplementary Figure 4G). In-gel fluorescence revealed an increased new/old ribosome ratio for the tagged 5K5R ribosomes relative to untagged ribosomes in the same cells; this was not observed in the wildtype tagged ribosomes (Figure 3F, Supplementary Figure 4H). This demonstrates that the increased ribosome degradation is specific to 60S ribosomes that fail to be UFMylated, and not an indirect effect of UFMylation loss altering cell state.

Given that the 60S subunit is the direct target of UFMylation and is degraded in response to lack of UFMylation, it is surprising that the 40S subunit was degraded more than the 60S (Figure 2C, 3B). We wondered if this small subunit response might be due to a lack of “free” cytosolic large subunits. Recent reports suggest that failures at initiation, including those caused by reductions in large subunit levels, cause 40S degradation via a pathway that depends on multiple mono-ubiquitinations by the E3 ligase RNF10, which are recognized by RIOK3 to commit the small subunit to degradation (Figure 3G).^35,44,88^ To test if this feedback might be the source of our small subunit degradation, we used RiboHalo flow cytometry to examine 40S and 60S subunit turnover when we knock down UBA5 (the UFMylation E1 enzyme), RIOK3, or both. Knockdown of RIOK3 had limited effects on old 40S or 60S subunit levels, although it mildly perturbed new levels of both (Figure 3H, Supplementary Figure 4I). As already observed, UBA5 knockdown led to reductions in old and new ribosomes of both subunits. Knocking down RIOK3 in the context of UBA5 knockdown completely rescued 40S subunit levels, with limited rescue of 60S. This is consistent with 40S degradation being a feedback response to large subunit loss via the RNF10-RIOK3 pathway. The degradation of the 60S subunit was not clearly due to a previously described pathway. We thus focused our further efforts on characterizing this large subunit degradation.

### Contact inhibition is required to promote large subunit degradation upon UFMylation loss

UFMylation loss had limited effects on ribosome levels in growing RPE1s compared to their quiescent counterparts (Figure 4A, Supplementary Figure 5A); no large subunit reduction was observed, and the magnitude of small subunit change decreased. We tested how contact-inhibition and serum reduction each contributed to this difference in degradation. Large subunit degradation upon UFMylation loss was independent of serum reduction; however, growing cells at lower confluence, even under reduced serum, eliminated the 60S subunit degradation phenotype (Figure 4B). This suggested that contact inhibition was required for large subunit degradation. Examining our genetic screens, we noted that knockdown of upstream members of the Hippo pathway were among the strongest pro-growth signals in our quiescent cell screens (Supplementary Figure 5B). We wondered if this pathway, which senses cell-cell contact and drives growth arrest by sequestering the pro-growth transcription factor YAP in the cytoplasm (Figure 4C),^89–91^ might be a required signal for large subunit degradation. To test this, we mimicked Hippo pathway activity by adding the YAP inhibitor GNE-7883,^92^ which disrupts its interaction with TEAD transcriptional coactivators. Adding this drug to nonconfluent cells rescued the degradation of large subunits upon UFMylation loss, confirming that Hippo pathway activity is required (Figure 4D).

**Figure 4:**
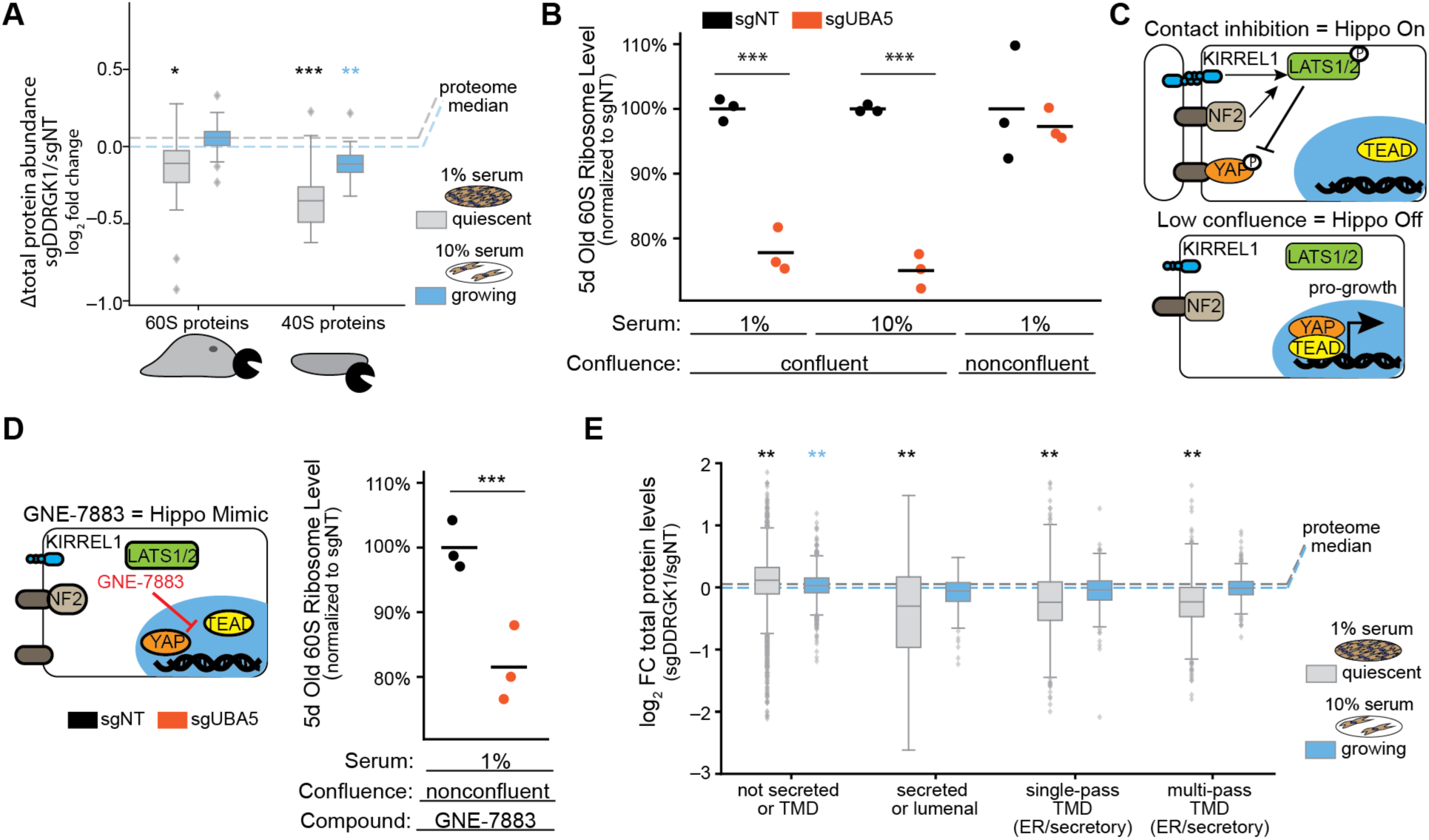
Large subunit degradation and reduced ER protein synthesis on UFMylation loss are dependent on contact inhibition A) Total protein level change differences for 60S and 40S subunit proteins upon DDRGK1 knockdown. *: p < 0.05, **: p < 0.01, ***: p < 0.001, Mann-Whitney U-test versus total proteome, Benjamini-Hochberg adjusted p-values. Dashed lines: median log_2_ fold change for quiescent (grey) and growing (blue) proteomes. B) Differences in old large subunit levels from a 5 day RiboHalo experiment (normalized per-condition to average of sgNT). ***: p < 0.001, Welch’s t-test. All points median fluorescence intensity from flow cytometry; horizontal line shows average. Serum and confluence as indicated. C) Schematic of Hippo pathway and confluence dependence. D) GNE-7883 mimics Hippo pathway activity at low confluence. Schematic and flow cytometry data as in Panel B. E) Changes in ER translocon-dependent protein levels upon DDRGK1 knockdown, in quiescent or growing conditions. Dashed lines and asterisks as in panel A.

Proteome-level changes upon UFMylation loss in quiescent revealed a broad reduction in proteins of the ER, Golgi apparatus, and plasma membrane relative to the rest of the proteome (Supplementary Figure 5B), all compartments dependent on protein biogenesis at the ER. This effect was specific to quiescent cells. As UFMylation is thought to play a role in ribosome release from the ER translocon, we wondered if this failure to resolve 60S-translocon complexes may impair subsequent ER protein synthesis. Consistent with this, we observed that loss of UFMylation led to reduction of soluble ER-translated proteins as well as transmembrane proteins in quiescent cells, but not their growing counterparts (Figure 4E); this trend is not observed for mitochondrial proteins, and tail-anchored proteins that are not translocon-dependent (Supplementary Figure S5C). Consistent with different responses to UFMylation loss between growing and quiescent cells, quiescent cells upregulate the translocon far less than growing cells upon UFMylation loss, and do not appear to upregulate ER-resident UPR markers such as the chaperone BiP, protein disulfide isomerase, or calreticulin (Supplementary Figures S5D–E).^93,94^ Taken together, this data suggests contact-inhibited cells face deficits in ER protein levels and degrade large subunits that fail to be UFMylated, while growing cells escape this level of ER translation impairment, potentially by upregulating the ER translocon core.

### Large subunits that fail to release from the ER are degraded

We hypothesized that failure to release the 60S ribosome from the ER translocon is driving large subunit degradation. Sucrose gradients from confluent cells showed that UBA5 knockdown led to reduced translation and an accumulation of non-translating 60S subunits (Figure 5A). To test if these non-translating large subunits might be failing to release post-termination, we turned to single-molecule tracking via microscopy (Figure 5B). By imaging ribosomes in confluent cells with highly inclined and laminated optical sheet (HILO) microscopy at short (15 ms) intervals,^95,96^ we were able to calculate diffusion rates of HaloTagged ribosomes using a statistical inference approach designed to account for short trajectory lengths and rapid defocalization.^95^ We observed that long-term UBA5 knockdown led to a notable shift in the relative diffusion rate occupancy of the 60S ribosomes, but muted effects on 40S diffusion (Figure 5C, Supplementary Figure 6A–B). We binned diffusion rates into five fractions, manually chosen based on troughs in probability density for 60S diffusion across experiments, and statistically compared the difference in fraction of ribosomes that had estimated diffusion rates in a given bin upon UBA5 knockdown. The large subunit showed reduced occupancy in a high-mobility state (bin IV) upon UFMylation loss and showed increased 60S ribosomes in lower mobility states (bins II and III, Figure 5C, Supplementary Figure 6C). Ribosome runoff using the translation initiation inhibitor harringtonine (which allows translation elongation and termination but prevents initiation, causing accumulation as free subunits and monosomes)^97,98^ and trapping experiments using emetine (which should irreversibly trap ribosomes in elongation phase, stabilizing polysomes and depleting free subunits)^98–100^ allowed state assignment for these diffusion “bins” as translation-dependent or free subunits (Supplementary Figures 6D–E). These results confirmed prior literature,^101–105^ that the two lowest-mobility bins are actively translating polysomes, while bin IV is likely free 60S subunits; we infer the highest-mobility bin is free ribosomal protein. Bin III, where 60S ribosomes accumulate upon UFMylation loss, was ambiguous with respect to ribosome inhibitor experiments, potentially indicating a mixture of species. These bin assignments suggested that our in-cell diffusion measurements contradicted our polysome analysis; while sucrose gradients showed an increase in free large subunit upon UFMylation loss, single-molecule tracking suggested a decrease.

**Figure 5:**
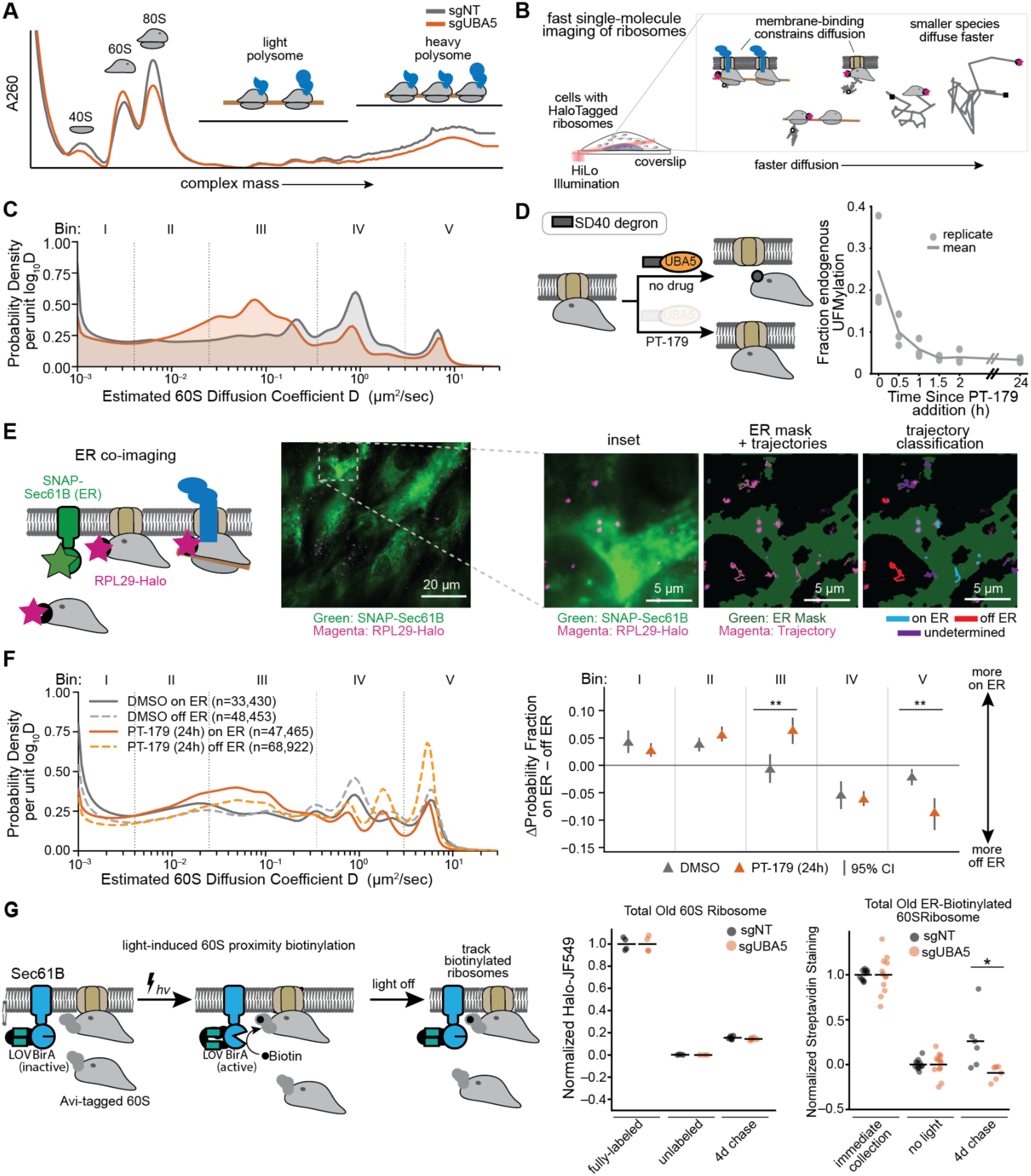
Ribosomal large subunits that fail to release from the ER are degraded faster A) Sucrose gradient comparing UBA5 knockdown to control. Symbols above represent molecular species. B) Schematic of single-molecule experiment. HILO microscopy was used to image individual ribosomes, and computational tracking allowed extraction of population diffusion rates. Inset shows example molecular species with different diffusion rates. C) Estimated diffusion distribution for RPL29-HaloTag. Numbers in legend are the number of subtrajectories analyzed from n=10 fields of view (FOV) per genetic condition. Vertical lines and bin labels designate regions for statistical comparison. Y-axis is posterior probability per unit log_10_D. D) Left: schematic of degron-inducible UFMylation reduction upon addition of degrader compound PT-179. Right: graph of RPL26-UFM1 loss as quantified by immunoblotting, compared to cells without manipulated UFMylation. E) Schematic and representative images of ER co-imaging. SNAP-Sec61ꞵ stains ER in a different channel from ribosomes. Image shows max intensity projection of 8 frames of ribosome imaging (magenta) with median projection of ER imaging (green). Inset shows boxed region, calculated ER mask, and trajectories before (middle) and after (right) categorization. Trajectories not enriched on or off mask are labeled “undetermined” and excluded from localization analysis. F) Estimated population diffusion rates for RPL29-HaloTag that are enriched on or off ER mask, before or after 24 hours PT-179 treatment to degrade UBA5. Numbers beside condition represent the number of subtrajectories analyzed (10 FOV DMSO, 14 FOV PT-179). Right: quantification of diffusion bins; vertical bars are 95% confidence intervals. ***: p < 0.002 (null minimum from FOV resampling). G) Schematic and data from optogenetic proximity labeling of ER ribosomes. LOV-BirA tethered to Sec61ꞵ biotinylates RPL29-Halo-AviTag 60S ribosomes at ER when light is on. Right: flow cytometry data showing HaloTag (total 60S) labeling and biotin staining. Fully-labeled samples were collected immediately after labeling, unlabeled collected without HaloTag dye addition, and 4d chase collected 4 days after HaloTag and Biotin labeling and chasing with nonfluorescent blocker. *: p<0.05, Welch’s t-test.

We hypothesized that the discordance in free large subunit levels between sucrose gradient and live-cell diffusion measurements is due to the emergence of an ER-bound large subunit population that fails to release from the ER translocon but would be solubilized by detergent during cell lysis for polysome preparation. This population would have lower mobility and may be part of the increased occupancy in diffusion bin III upon UBA5 knockdown. We additionally wanted to rule out that large subunit accumulation was a consequence of increased ER volume and translocon abundance, known feedback responses to a lack of UFMylation.^84,106^ To address this ER-expansion concern, we generated a degron-tagged copy of UBA5 which we used to replace endogenous UBA5 after CRISPRi knockdown (Figure 5D, Supplementary Figures 6F–G).^107^ While this system showed reduced overall UFMylation levels (Supplementary Figure 6F), it provided temporal control, allowing ∼4-fold reduction in UFMylation levels within two hours upon addition of degrader compound PT-179 (Figure 5D, Supplementary Figure 6F). We reasoned that this rapid temporal control could allow us to examine the effects of UFMylation loss on ribosome mobility before long-term cellular adaptation. At longer timepoints, these degron reporter lines showed increased ribosome degradation upon UFMylation loss for both 40S and 60S ribosomes (Supplementary Figure 6G).

In this rapid-UFMylation-depletion system, we introduced an orthogonal self-labeling protein, SNAP-tag,^108^ onto Sec61ꞵ, a tail-anchored ER protein that acts as a general ER marker.^109,110^ We imaged this ER marker for 100 frames prior to imaging 60S ribosomes for single-particle tracking and constructed a mask to classify trajectories as “on-ER” or “off-ER” (Figure 5E, Supplementary Figure 6H). The area of this mask did not increase after 24 hours of PT-179 addition, suggesting limited ER expansion (Supplementary Figures I). Compared to long-term UBA5 knockdown, short-term UFMylation loss showed less global differences in 60S diffusion between PT-179 treated and DMSO controls (Supplementary Figure 6J). However, comparing between the “on-ER” and “off-ER” populations, we found that diffusion bin III accumulated in the “on-ER” population in PT-179 treatment, but showed no difference in the untreated control (Figure 5F). This provides direct support for ourhypothesis that the 60S is retained on the ER after termination, and this accumulation is induced upon UFMylation loss as a retained free 60S.

Having established that 60S ribosomes accumulate on the ER upon UFMylation loss, we wanted to determine whether the degraded population of 60S ribosomes is this ER-bound subset. In order to test this hypothesis, we adapted our recently published protocol for light-inducible proximity biotinylation of ribosomes at the ER (Figure 5G).^111^ This system activates a biotinylating enzyme, LOV-BirA, using blue light: when active, the enzyme specifically attaches biotin to proteins bearing the Avi-tag peptide tag that come into physical contact.^112^ Turning off the blue light inactivates LOV-BirA, ceasing the biotinylation of new molecules brought into proximity but allowing us to track the fate of those molecules biotinylated during the labeling period. We used a previously characterized ER-bound LOV-BirA alongside overexpressing RPL29 tagged with both HaloTag and Avi-tag, allowing us to track both total 60S and ER-bound 60S fates from the same set of ribosomes. Introducing these constructs alongside either UBA5 knockdown or control CRISPRi guides allowed us to track effects of UFMylation loss on total and ER-localized 60S degradation. No difference was observed between 60S HaloTag levels at a 4 day timepoint but we saw clear differences in the ER-labeled subset, with biotin signal having returned to the unlabeled baseline in the UBA5 knockdown condition, but not in the control cells. Taken together, the combination of sucrose gradient analysis, live-cell biophysics, and proximity labeling establish that upon UFMylation loss, 60S ribosomes accumulate at the ER outside polysomes and that these ER-accumulated ribosomes are the subset that are rapidly degraded. To our knowledge, this degradation of ER-bound 60S ribosomes has not previously been reported-we term this phenomenon “<u>T</u>ranslocon-associated <u>L</u>arge <u>S</u>ubunit <u>D</u>egradation”, or TLSD.

### Translocon-associated large subunit degradation is translation-dependent and autophagy-independent

We sought to better characterize TLSD via a CRISPRi suppressor screen (Figure 6A).^113^ We used the same multi-sort screening strategy described above to measure the genetic interactions between UFMylation knockdown and knockdown of other pathways; by measuring the change in degradation score, we could classify genes as acting on degradation via related or unrelated pathways based upon degree of epistasis (Figure 6A, right).^114,115^ We used the same sublibrary as initially used to determine degradation and synthesis scores with a fixed second guide in a dual-guide vector targeting UBA5 knockdown (Figure 6B, Supplementary Figures 7A–B, Supplementary Tables S10–S11), allowing direct comparison to our earlier sublibrary screens (Figure 2).

**Figure 6:**
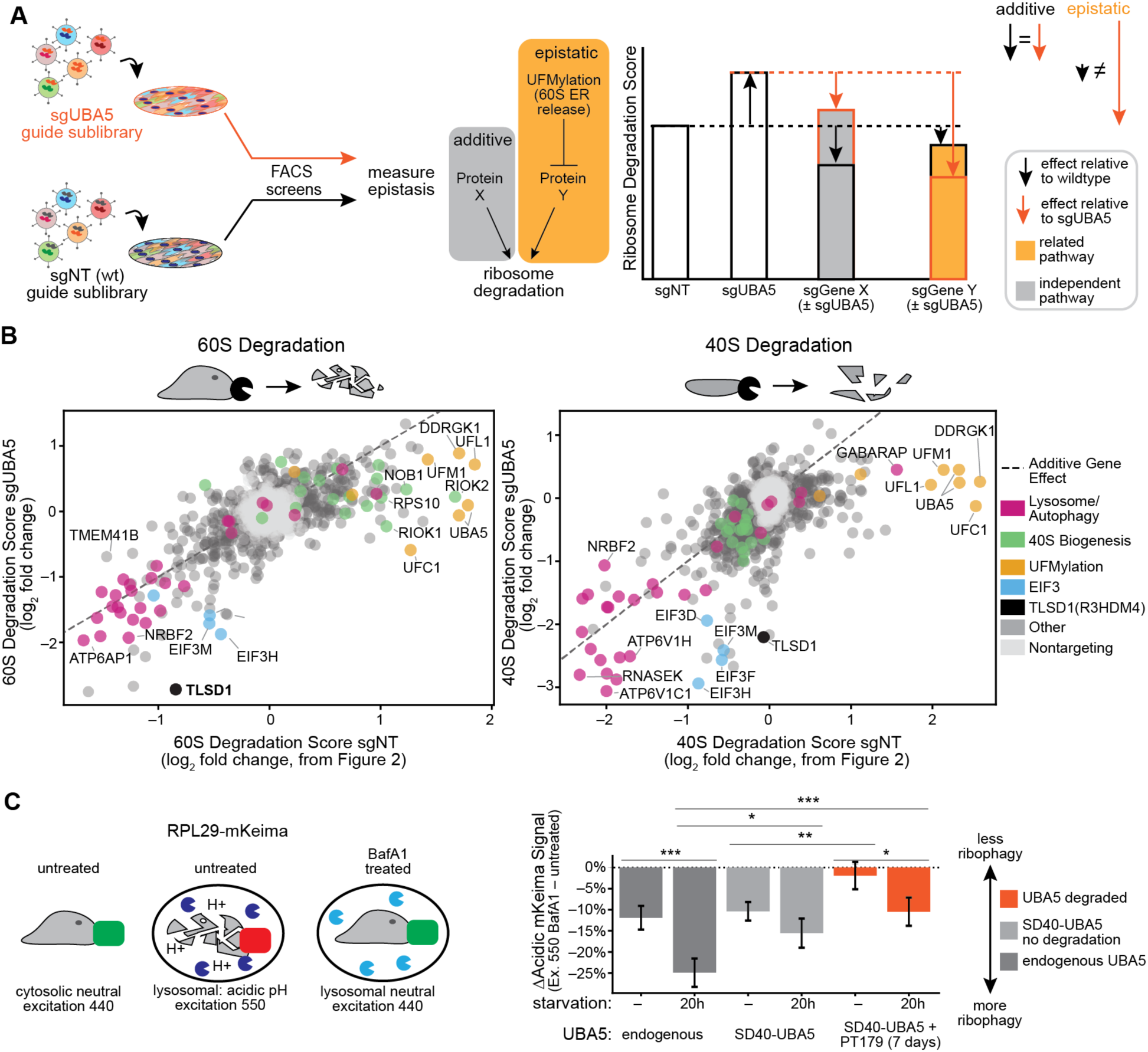
Suppressor screens identify mechanistic aspects of translocon-associated large subunit degradation A) Schematic of dual-guide suppressor screening. In one library, a fixed control guide (sgNT) was included alongside sublibrary (data shown in Figure 2); in the second, a fixed sgUBA5 guide was used, and both libraries subjected to multi-sort strategy to extract degradation and synthesis scores. Right shows epistasis logic used to identify TLSD-related genes. B) Suppressor screen results for 60S (left) and 40S (right) degradation, showing degradation score relative to control (x-axis) or fixed UBA5 knockdown (y-axis). Dashed line shows additive effects on degradation score between screens. Colored based on gene categories. C) Schematic (left) and median flow cytometry data (right) for Bafilomycin A1-dependent acidic RPL29-mKeima signal. Data shows the difference in acidic signal upon overnight treatment with 10 nM Bafilomycin A1 (BafA1) under starvation, UFM1 degradation for 7 days, or both, as percentage change between BafA1 treated and untreated. Starvation removed serum or amino acids as a positive control for autophagy induction for indicated time. Bars are standard deviation from n=6 treated and untreated wells each. *: p<0.05, **: p < 0.01, ***: p < 0.001, Benjamini-Hochberg corrected Welch’s t-test.

Degradation and synthesis score epistasis revealed several aspects of TLSD. First, confirming the effectiveness of the epistasis strategy, we saw reductions in degradation score for UFMylation-pathway gene knockdown on top of UBA5 knockdown (Figure 6B), consistent with the TLSD already being saturated by UBA5 knockdown. Intriguingly, the 60S degradation seen upon late 40S maturation defects was also suppressed, suggesting a shared degradation pathway promoted via two genetic pathway disruptions. Synthesis scores showed limited epistatic interactions (Supplementary Figure 7B–C), although genes that increased 40S synthesis had blunted responses. Knockdown of components of EIF3, a complex required for initiation of translation,^116^ suppressed degradation of both subunits when UFMylation was absent but had muted effects in its presence. We took this as evidence that TLSD was translation dependent. In contrast to the epistasis upon EIF3 knockdown, we noted that although autophagy and lysosomal ATPase knockdown slowed degradation in both screens, they showed no genetic interaction with UBA5 knockdown, having near-identical degradation scores. We confirmed this result using overexpression of an RPL29-mKeima reporter, a pH-sensitive fluorophore used to measure autophagic flux.^117^ Comparing flow cytometry-derived fluorescence before and after treatment with Bafilomycin A1 (BafA1)^118^ showed that UBA5 degradation led to a decreased baseline ribophagy (Figure 6C, Supplementary Figure 7D), consistent with prior reports suggesting that UFMylation knockdown suppresses rather than promotes autophagy.^71^

### Gene of unknown function TLSD1 plays key role in translocon-associated large subunit degradation

The most intriguing finding from our suppressor screening experiment was the discovery that knockdown of R3H-domain containing 4 (R3HDM4), a gene of unknown molecular function, strongly suppressed degradation of the large subunit(Figure 6B). We propose re-naming this gene for its apparent role in TLSD as TLSD1. TLSD1 is among the least-well annotated genes in the human genome,^119^ but is broadly conserved in vertebrates;^120^ the only clear domain is an R3H domain, which is known to engage in single-stranded nucleic acid binding (Supplementary Figure 8A).^121^ In our wildtype sublibrary screens, knockdown of TLSD1 mildly decreased 60S (but not 40S) new/old ratio (Figure 6B), and had a mildly growth-increasing effect (Supplementary Figures 7E). In the context of UFMylation knockdown, TLSD1 knockdown greatly decreased 60S degradation and it moderately reduced 40S degradation (Figure 6B). TLSD1 knockdown had the strongest pro-growth epistasis in our sublibrary, consistent with its loss preventing ribosome degradation and allowing higher translation rates (Supplementary Figure 7E–F). We confirmed the rescue of ribosome levels in RiboHalo flow cytometry assays (Figure 7A, Supplementary Figure 8B–D), and noted that 60S subunit levels were increased beyond control levels. The 60S accumulation appeared to be specific to old large subunit levels (Figure 7A, Supplementary Figure 8B), confirming the effect is due to degradation and not biogenesis.

**Figure 7:**
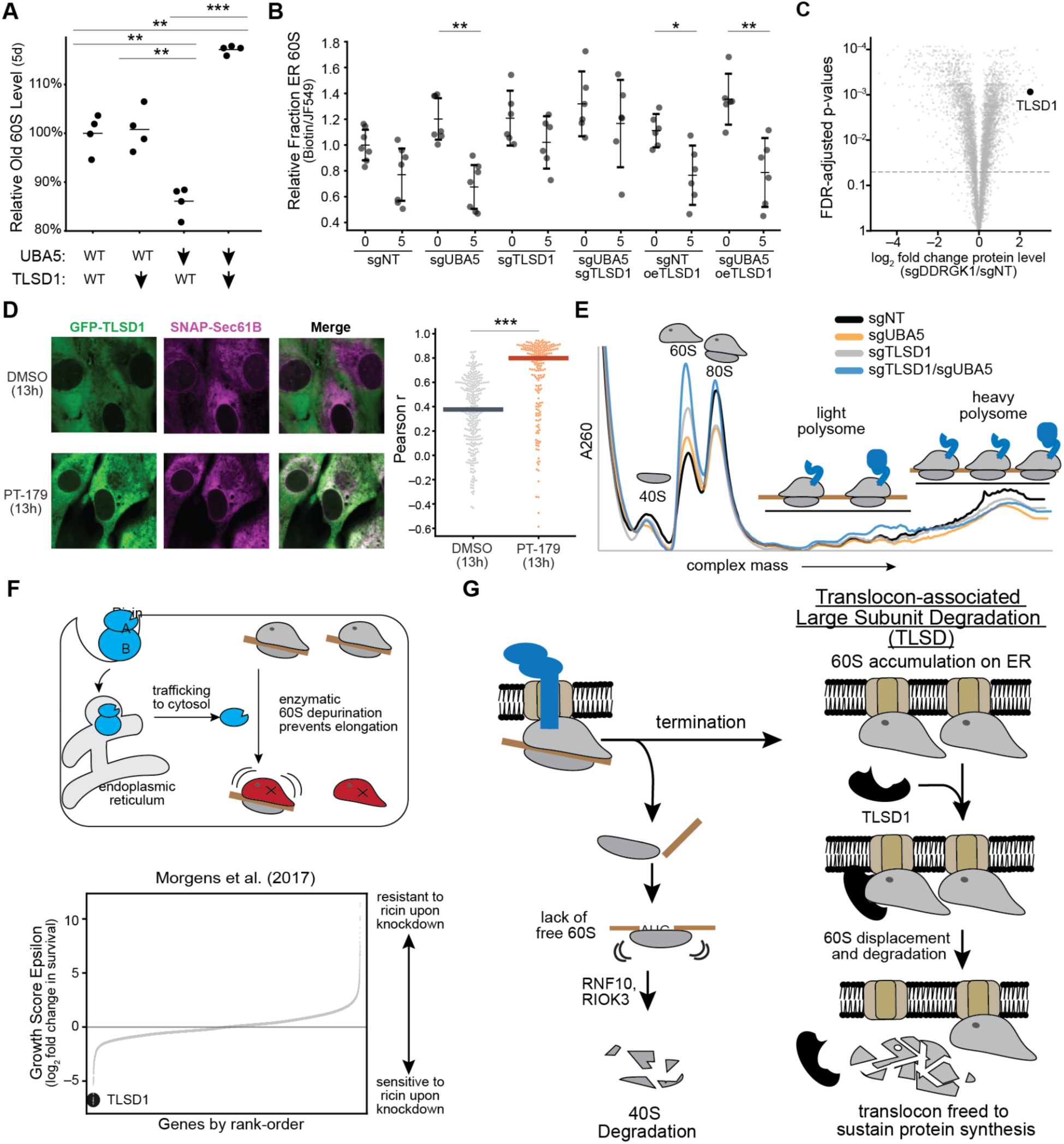
TLSD1 is required for degradation of ER-retained ribosomes A) Effect of UBA5 knockdown, TLSD1 knockdown, or both on >5-day old 60S ribosome levels by flow cytometry. Individual points represent median fluorescence, normalized to mean of sgNT. **: p < 0.01, ***: p < 0.001, Benjamini-Hochberg corrected Welch’s t-test. B) Relative fraction of ER-labeled ribosomes at 0 and 5 days from PAGE-based proximity labeling assay. Total ribosomes labeled with JF549-Halo, ER ribosomes biotinylated by LOV-BirA-Sec61ꞵ; samples collected immediately after labeling or 5 days later. Biotin signal (ER ribosomes) per JF549 (total ribosomes) normalized to sgNT at time=0. n=6 per condition, each dot is independent replicate. Vertical bars: 95% confidence interval. *: p<0.05, **: p<0.01, Welch’s t-test, Benjamini-Hochberg corrected. C) log_2_ fold change protein levels (day 8, light SILAC labeled) between sgDDRGK1 and sgNT in quiescence vs. Benjamini-Hochberg-adjusted p-value. TLSD1 highlighted. D) Live-cell confocal microscopy of GFP-TLSD1 and SNAP-Sec61ꞵ as ER marker in cells with SD40-UBA5. Left-inset showing each channel and overlay. Right-graph showing per-cell correlation coefficient between SNAP-Sec61ꞵ and GFP-TLSD1. Treatment for 13 hours with 20 µM PT-179. E) Sucrose gradient traces from different genetic conditions (sgNT and sgUBA5 as in Fig. 5A). Traces normalized by total A_260_. Cartoons represent molecular species. F) Left: schematic of ricin as poison that enters through endocytic pathway and damages 60S ribosome. Right: Waterfall plot of growth sensitization and resistance upon ricin treatment relative to control. Data from Morgens et al. 2017.^122^ G) Schematic of events upon loss of UFMylation, showing 40S degradation via RNF10-RIOK3 pathway, and TLSD-mediated degradation of 60S and preservation of ER translation.

We sought to further characterize the effects of manipulating TLSD1 on 60S degradation.

By combining LOV-BirA-Sec61ꞵ proximity-labeling system with RiboHalo labeling to track both total 60S ribosomes and ER-proximal ribosomes while manipulating UBA5 levels (CRISPRi knockdown) alongside TLSD1 levels (either CRISPRi knockdown or overexpression), we could determine relative rates of ER and total 60S ribosome degradation. In these assays, UBA5 loss caused an increase in both total 60S signal and ER-localized ribosome signal at early timepoints (Figure 7B, Supplementary Figure 8E–G). At 5 days after 60S labeling, cells with UBA5 knockdown showed accelerated degradation of ER ribosomes relative to total ribosomes (Figure 7B, Supplementary Figure 8F–G), and knockdown of TLSD1 prevented this degradation. TLSD1 overexpression was sufficient to accelerate ER 60S degradation in the absence of UBA5 knockdown, suggesting a basal role at the ER for TLSD1. While overexpressing TLSD1 alongside UBA5 knockdown led to lower total 60S levels (Supplementary Figure 8F), the fraction at the ER was high at early timepoints (Figure 7B, Supplementary Figure 8G). By 5 days, this increased ER-bound 60S fraction had been degraded and was at similar levels to cells that never had increased ER levels (Figure 7B, Supplementary Figure 8G). Taken together, these data suggest TLSD1 overexpression does not prevent ER ribosome accumulation, but degrades the ribosomes trapped on translocon, preventing them from persisting and decreasing the total large subunit pool. Importantly, in a recently-reported gel-based metric of autophagy based on the appearance of an autophagy-dependent HaloTag cleavage product,^123,124^ TLSD1 knockdown or overexpression had no effect on 60S ribophagy, and UBA5 knockdown suppressed ribophagy rather than increasing it (Supplementary Figures 8H–I), further confirming that TLSD does not accelerate ribosome degradation via enhanced ribophagy.

In examining TLSD1 further, we found it was at the right subcellular location, expressed at the right time, and its presence seemed to play a role in effective translation. TLSD1 is among the most strongly induced proteins upon DDRGK1 knockdown (Figure 7C), suggesting that failure to release the 60S ribosome from the ER increases TLSD1 levels. Consistent with TLSD1’s role at the ER, microscopy analysis of GFP-tagged TLSD1 showed a shift from diffuse cytoplasmic GFP signal to TLSD1 co-localization with an ER marker upon UBA5 degradation (Figure 7D). Sucrose gradients showed that TLSD1 knockdown elevated free 60S levels but had mild effects on polysome shape (Figure 7E). However, upon co-knockdown of UBA5, we saw further increases in 60S levels, and increased light polysomes without a rescue of heavy polysomes, suggesting that the 60S ribosomes that were no longer degraded upon TLSD1 loss were capable of initiation but did not support robust translation; this ability to initiate is consistent with the rescue of 40S levels by TLSD1 knockdown.

Finally, to help explore functional roles TLSD plays when UFMylation is not perturbed, we searched the literature for phenotypes for TLSD1 in prior CRISPR screening efforts.^125^ Intriguingly, the most prominent phenotype comes from re-analysis of growth screen data examining changes in survival upon ricin treatment in the leukemic cell line K562 (Figure 7F).^122^ Ricin is an enzymatic toxin that depurinates A4605 in the sarcin-ricin loop of the large subunit’s 28S rRNA, preventing elongation factor binding and ribosome translocation.^126,127^ Previous analysis of this screen focused on the known resistance mechanisms as positive controls.^128^ In our re-analysis, we find TLSD1 loss causes a profound and specific sensitization of cells to ricin (Supplementary Figure 7E). This is even more notable because TLSD1 loss mildly increases growth both in our screens and cancer cell lines broadly^129^ when a ribosome-damaging toxin is not present. This ricin sensitization would make sense if TLSD1 recognizes and commits non-functioning large subunits to degradation. By removing damaged large subunits, TLSD1 could enable the few non-damaged 60S ribosomes in cells that had been exposed to ricin to not be stymied by damaged ribosome “roadblocks” and engage in new protein production to support cell recovery. Taken together with our results in RPE1, we believe this supports a model of TLSD as a sensor for 60S damage, using ER release as a checkpoint to recognize problematic 60S ribosomes and commit them to degradation (Figure 7G).

## Discussion

Ribosomes are long lived structures^11–14,130^ that are susceptible to dysfunction,^18–20,127,131^ Despite decades of study dating back to the founding of molecular biology, their degradation remains poorly understood in mammals.^28^ This is particularly true for the large subunit, for which no selective degradation pathways for mature particles had previously been reported. Here, we developed a screening strategy to systematically map the genetic determinants of ribosome synthesis and degradation in quiescent human cells. This approach recovered previously known pathways of ribosome biogenesis and bulk turnover via autophagy; in addition, we uncovered two triggers for 60S ribosome degradation. The first was that 60S subunits are degraded in response to defects in late 40S biogenesis; the second was the discovery of translocon-associated large subunit degradation, or TLSD, which degrades 60S ribosomes that fail to release from the ER translocon in nondividing cells, which we followed up on in mechanistic detail.

Our findings support a model in which failure to release from the ER translocon targets 60S subunits for TLSD (Figure 7G). After translation termination at the ER, the 40S ribosome is released into the cytoplasm but the Sec61-60S ribosome complex remains stable. Release of the 40S subunit exposes the 60S subunit interface allowing recruitment of the UFMylation E3 ligase complex, which under normal circumstances transfers UFM1 onto the C-terminal tail of RPL26, driving a conformational shift that displaces the Sec61 translocon.^84,85^ In the absence of this modification, the 60S-translocon complex is stable;^87^ consistent with this, our single particle-tracking, proximity labeling, and sucrose gradient experiments demonstrate that large subunits accumulate at the ER as non-translating 60S ribosomes when UFMylation is lost (Figure 5, 7B, Supplementary Figure 8E–G). Structural snapshots of the UFMylation process suggest that post-termination 60S ribosomes are bound by the anti-association factor EIF6, preventing 40S binding to the subunit interface.^84,85^ This leaves a deficit of 60S ribosomes for subunit joining at translation initiation. Failed initiation leads to 40S ubiquitination by RNF10 and RIOK3-mediated degradation,^44^ preventing spurious leaky scanning and explaining why 40S ribosomes were sensitive to all perturbations that altered 60S levels (Figures 1F, 3H, 7G left branch, Supplementary Figure 3A). In quiescent cells, ER-retained 60S ribosomes are degraded in a TLSD1-depenent manner and TSLD1 is recruited from the cytoplasm to the blocked translocon upon 60S accumulation (Figures 7D, 7G right branch). TLSD1 levels increase in response to failure of 60S release (Figure 7C), and TLSD1 overexpression in a UFMylation-deficient background lowers total ribosome levels and accelerates their degradation at the ER, but does not reduce the total level of ER-bound 60S ribosomes, suggesting that it promotes 60S release from the translocon into a degradation pathway but does not interfere with upstream binding and accumulation. While the exact molecular mechanism of degradation remains to be determined, our proteomics shows that the translocon is not co-degraded (Supplementary Figure 4F), and multiple assays (Figure 6C, Supplementary Figure 8H–I) show this degradation is not via autophagy, suggesting cytoplasmic degradation that would require disassembly of the ribosome, at which point the individual ribosomal proteins and rRNA would be unstable and subject to proteasomal and nucleolytic degradation.^59,60,132^

It is unclear why post-termination accumulation at the ER is one of the few circumstances in which cells choose to degrade 60S ribosomes. In the absence of TLSD1, 40S levels are somewhat rescued, as are polysomes (Figures 6B, 7E, Supplementary Figure 8B–D). This suggests that the accumulated 60S ribosomes are able to initiate, if poorly, given the preponderance of free 60S and the lower number of ribosomes per mRNA observed in polysome traces. We hypothesize that in the absence of UFMylation and TLSD as a backup system, these accumulated 60S remain translocon-associated, and, if not bound by the UFMylation E3 machinery, are competent for EIF6 eviction and re-initiation while on the translocon.^133^ This would allow continued ER protein synthesis, potentially explaining the limited toxicity of losing UFMylation or both UFMylation and TLSD. In this instance, translocon binding would no longer be gated by SRP handoff, reducing efficiency of mRNA targeting during the pioneer round of translation, due to limited free translocon capacity for new ribosomes. Furthermore, cytosolic proteins would initiate on these 60S-translocon complexes. While the gating mechanism on Sec61 itself would limit mistranslation into the ER lumen,^134^ the mistethering of non-secretory protein synthesis to the translocon would limit accessibility of co-translational protein folding factors and quality control machinery to the ribosome exit tunnel, reducing efficiency of nascent protein folding and impairing clearance of peptides resulting from stalled protein synthesis.^75,135^ This would also further reduce the percentage of translocons engaged in translation of messages meant for the ER. It has long been proposed that a broad range of proteins are synthesized at the ER surface beyond secretory and membrane proteins;^136^ however, the existence of a pathway to degrade 60S ribosomes that fail to release from the translocon argues that remaining directly tethered is deleterious enough to provide an evolutionary pressure for their ribosome clearance.

One surprising aspect of our findings is the cell-state dependence of TLSD: 60S ribosome release failure leads to large subunit degradation only in contact-inhibited RPE1s. In the growing state, loss of UFMylation leads to significant upregulation of the ER translocon, and cells show neither global reductions in secretory protein synthesis or 60S protein levels, even though 40S levels still drop, suggesting some 60S ribosomes are still sequestered from initiation. During contact inhibition, translocon upregulation is muted but there are global secretory protein defects (Figure 4, Supplementary Figure 5); likely to prevent further defects, cells degrade these trapped 60S via TLSD. Intriguingly, knockdown of TLSD1 reduces fitness in CRISPR screens in iPSC-derived glutamatergic neurons,^137^ suggesting TLSD may play protective roles in other nondividing contexts.

We observed a second trigger of 60S degradation in our data: cytosolic 40S maturation defects. In yeast, 40S biogenesis involves the engagement of late 40S intermediates with mature 60S in a translation-like cycle that acts as a quality control checkpoint; failure to dissociate these 80S-like intermediates leads to degradation of both immature 40S subunits as well as bound mature 60S,^10,138^ a phenotype strikingly similar to that seen in our data. However, in humans, structural evidence has argued against this 60S-joining being a conserved process. ^80,139^ Our genetic results suggest that while the specific 80S-like intermediates may not be conserved from yeast to humans, there is a common link between late 40S maturation and the mature 60S that will require further mechanistic elucidation. Intriguingly, our genetic suppressor screen found that accelerated 60S degradation due to late 40S biogenesis knockdown was suppressed in cells with UBA5 knockdown, suggesting a role for TLSD in communicating defects in 40S biogenesis to the mature 60S ribosome (Figure 6B).

The broader physiological importance of TLSD remains to be fully elucidated; however, examination of TLSD1 from the lens of evolutionary conservation, prior human genetics, and CRISPR screens all point to important roles for this pathway. TLSD1 is conserved in vertebrates,^119^ and while molecular functions have not been previously ascribed, one report suggests it is associated with poor prognosis in kidney renal cell carcinoma.^140^ GWAS studies have linked TLSD1 expression to late erythropoiesis,^141^ which is one of the few physiological occasions in which a cell degrades its ribosomes, and impairment in ribosome elimination is known to alter this process;^142^ a role for TLSD in promoting ribosome elimination would align with the stage of erythropoiesis that appears to be altered. As noted above, CRISPR screening has shown sensitization to ricin, which targets the 60S, upon TLSD1 loss^122^ and loss of TLSD1 is detrimental to neurons, which are particularly sensitive to defects in translation, with mutations in translation quality-control or proteostasis machinery frequently causing neuronal or CNS phenotypes.^143–147^ We posit these results can together be explained if TLSD acts as a degradative quality-control checkpoint for the 60S ribosome, using release from the translocon as a mechanism to recognize dysfunction in order to degrade damaged subunits rather than recycling them. In rapidly dividing cells, increased biosynthetic capacity from larger numbers of ribosomes is a major driver of growth;^76^ although impaired ribosomes may generate stress, they will be diluted by the pool of new ribosomes. However, nondividing cells lack access to dilution and cells such as neurons would have higher sensitivity to defects in faulty ribosome removal. Combined with the cell-state-dependence of 60S removal, these results point to TLSD as a key pathway for those cells that need

60S quality control, but one kept under strict control to prevent spurious ribosome degradation from limiting cell growth.

Cell-state-dependent differences in proteostasis have emerged as an important theme in recent years, and have important implications for understanding pathogenesis in diseases of aging and protein misfolding, such as Alzheimer’s disease, Parkinson’s disease, and other neurodegenerative diseases.^148–150^ Notably, these diseases often see early dysfunction in ribosomes;^27,151^ however, while alterations in overall translation and autophagy rates have been heavily explored in these contexts,^5,152^ there has been little exploration of pathways of ribosome turnover. Our work reveals a new pathway of ribosome degradation that adds to the list of cell-state-dependent quality control pathways. Further extending the strategies we use here in other cell models would not only deepen our understanding of fundamental biology but may provide actionable insights into disease pathogenesis.

### Limitations

By using a genetic knockdown approach, we are able to discover genes whose absence alters degradation rate; however, if a gene is not expressed or active at baseline, we cannot observe a reduction in degradation. This suggests that our screening approach provides a genetic map of active degradation pathways in the particular cell state used for screening-if a damaging event does not occur in this state, it will not be picked up by the genetic approach. Our follow-up focused on the use of a sublibrary derived from the primary screen; this allowed us to discover TLSD1, but there may be other TLSD genes that did not sufficiently contribute at baseline to be included in this sublibrary.

## Resource Availability

### Lead Contact

Further information and requests for resources and reagents should be directed to and will be fulfilled by the lead contact, Jonathan S. Weissman.

### Materials Availability

All unique reagents generated in this study are available upon request; plasmids will be made publicly available through Addgene upon publication.

### Data and Code Availability

All code used for analysis will be publicly available from GitHub upon publication and is available upon request. CRISPRi screen sequencing data will be deposited at GEO and proteomics data will be deposited at ProteomeXchange via PRIDE and publicly available after publication; imaging data, due to size constraints, will be available upon request, while trajectory data will be publicly available at Zenodo and is available upon request. Plasmids and cell lines generated in this study are available from the lead contact upon request, and plasmids will be deposited on Addgene. Lead contact: Jonathan S. Weissman

## Acknowledgements

We thank A. Guna, D. Yang, L.W. Koblan, T. Bertozzi, K. Smolyar, P. Zhang, G. Muthukumar, R.S. Saunders, M. Brolosy, K. Yost, V. Baumann, and members of the Weissman Laboratory for reagents, advice, and technical assistance. We also thank J. Paulo and E.L. Huttlin from Harvard Medical School for proteomics acquisition and analysis assistance. We also thank P. Autissier, A. Rathee, and K. Goto-Hardy from the Flow Cytometry Core Facility at Whitehead Institute for FACS sorting. We thank M. Mazzocca and P. Zhang for microscopy assistance. We also thank C. McCabe for assistance with sequencing. This work was supported by the Helen Hay Whitney Foundation Fellowship (Z.G.L. and K.L.H.), the Damon Runyon Cancer Research Foundation Fellowship (DRG-2411-20 to J.L.), and NIH (F32GM150241 and K99GM166795 to J.J.). J.S.W. is an HHMI investigator.

## Author Contributions

Z.G.L and J.S.W conceived the project and designed the experiments. K.L.H. carried out proteomics sample preparation and analysis in the laboratory of J.W.H. Z.G.L. and G.S. designed the single-molecule experiments, carried out in the laboratory of A.S.H. J.J. cloned genome-wide library; Z.G.L. and S.K. cloned sublibraries. J.J. and Z.G.L. designed CRISPR screen analysis. Z.G.L., S.K., C.J.Z., and J.H. performed cloning and carried out targeted turnover assays. Z.G.L., J.H., and J.L. designed and carried out the proximity labeling experiments. Z.G.L., Y.H.C-N., and S.K. carried out CRISPR screening experiments.

## AI Disclosure

Claude Opus models 4–5 were used in the generation of analysis and plotting code during this work; all workflows were generated as directed by humans and code verified for accuracy. No writing was directly generated by AI; AI was used as an aid in the editing process for grammar and proofreading.

## Supplementary Figures

**Figure S1:**
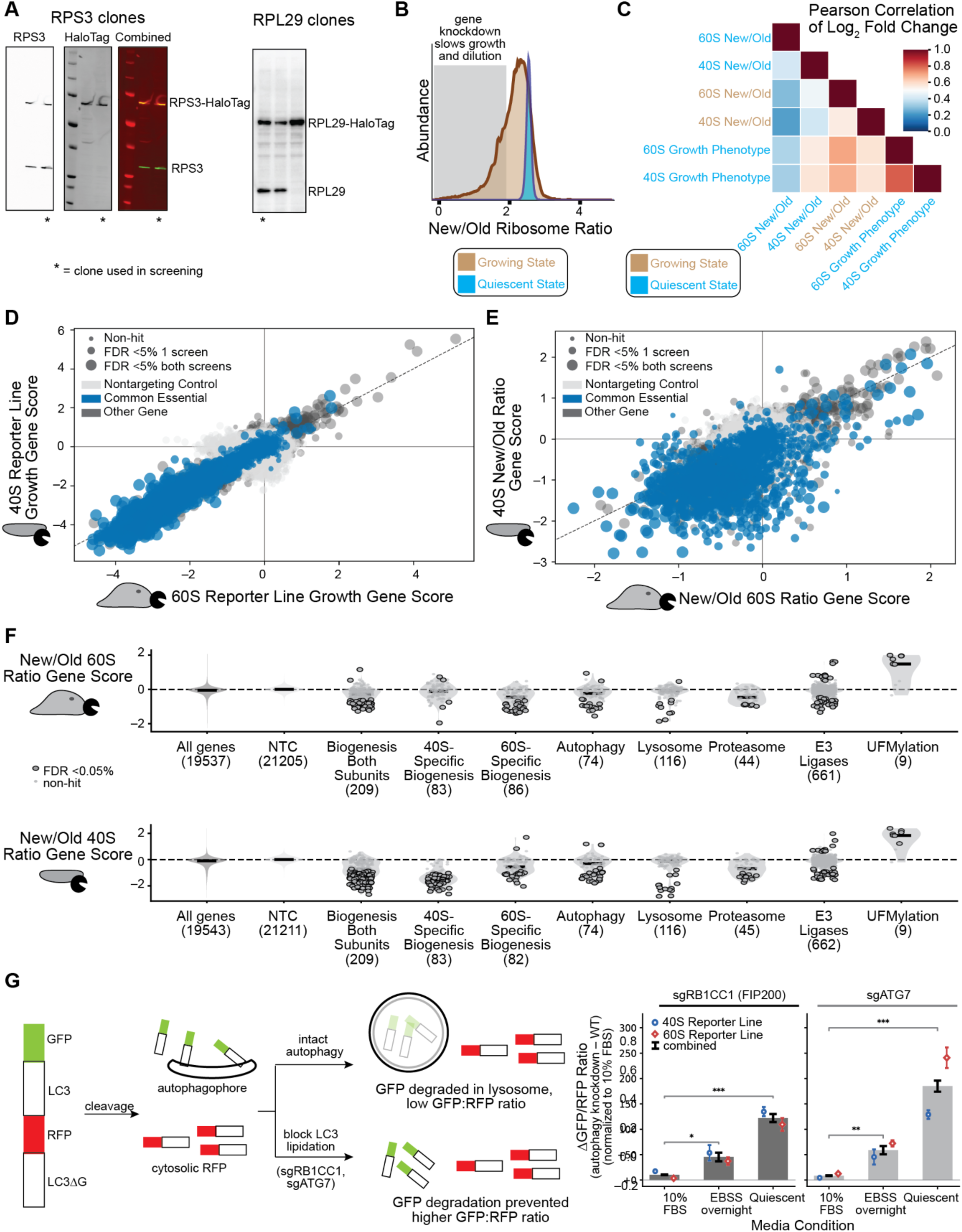
Genomewide HaloTag turnover screening in quiescent RPE1s allows systematic measurement of genetic determinants of ribosome turnover (related to Figure 1) A) Immunoblots showing clones isolated from endogenous genome editing of RPS3 (left) and RPL29 (right) with HaloTag7. Heterozygous expressing clones were chosen (asterisks) to move forward with to be consistent between subunits. B) Histograms of new/old ribosome ratio from cells infected with genome-wide library, from 3-day growing cell turnover experiment and 9 day quiescent cell turnover experiment C) Pearson correlation of gene score (log_2_ fold change) for cells sorted on new/old ribosome ratio under quiescent or growing conditions, or evaluated for change in guide abundance between early timepoints and screen endpoint in quiescent cells (“Growth phenotype”) D) Quiescent state growth gene scores (log_2_ fold change) for each reporter cell line, highlighting common essentials from DepMap. Small subunit on Y-axis, large subunit on X-axis. E) Quiescent state new/old ratio gene score (log_2_ fold enrichment) for small subunit (Y-axis) and large subunit (X-axis), with DepMap common essentials highlighted. F) Gene score for new/old ribosome ratio for specific categories of genes by subunit. Each individual point is a gene, outlined points meet 5% FDR cutoffs in Mann-Whitney test against non-targeting controls, and overall distribution is represented by grey “violin” in background. Distribution-only shown for all genes and non-targeting controls. Numbers in parentheses indicate number of genes in category as plotted. G) Dual-color bulk autophagy reporter. The reporter cotranslationally produces GFP-tagged LC3 and RFP-tagged LC3ΔG, which is not incorporated into autophagosomes. During autophagy, GFP is trafficked to lysosome and reduced. Knockdown of ATG7 or FIP200 blocks canonical macroautophagy, and thus knockdown would be expected to increase GFP:RFP ratio. Data at right is change in GFP:RFP ratio upon gene knockdown under different media conditions. Each was performed in triplicate per reporter cell lines, with bars representing total mean difference between gene knockdown and wildtype control, with bars showing combined standard error, calculated per cell line (and combined).

**Figure S2:**
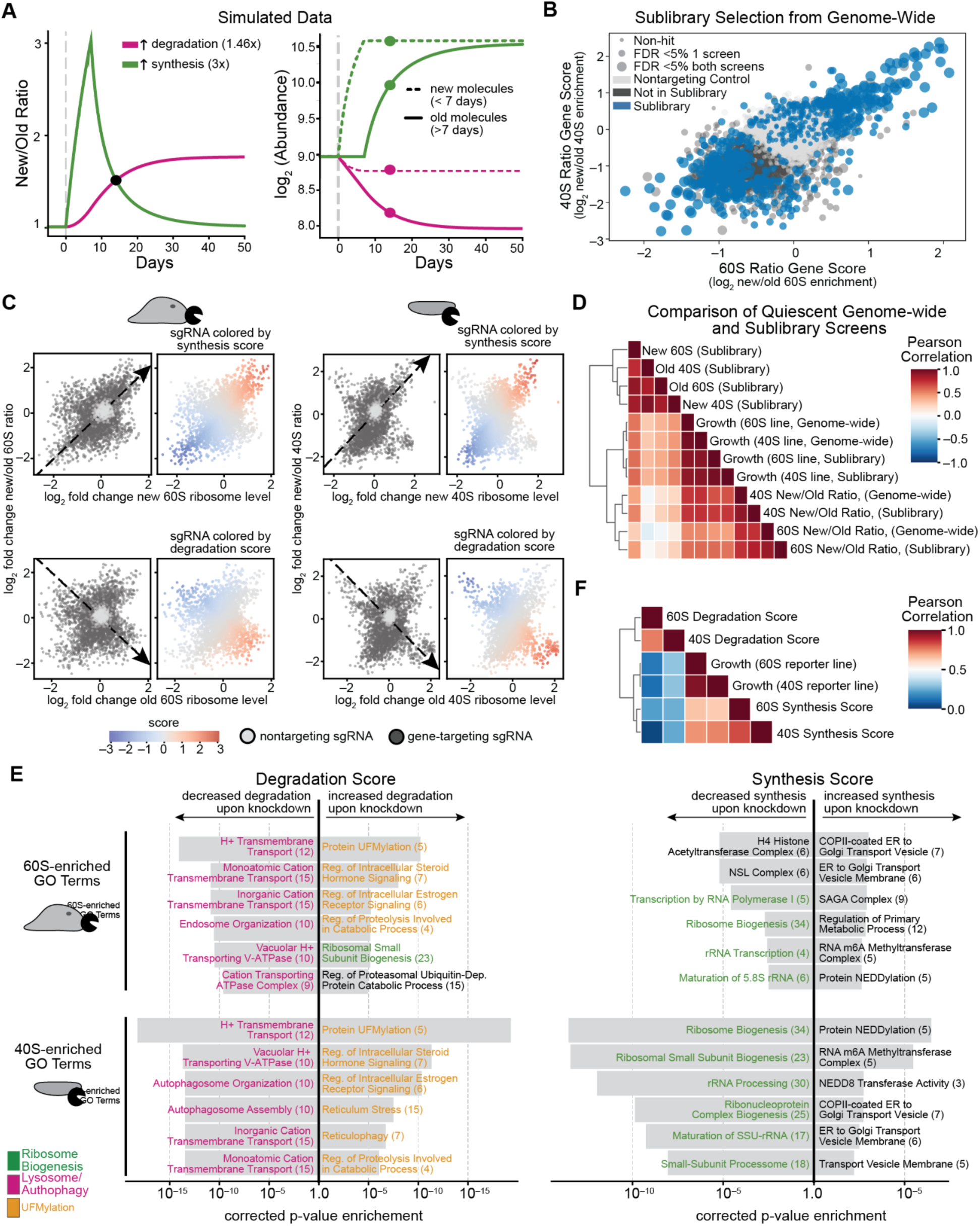
Sublibrary screening allows classification of synthesis and degradation (Related to Figure 2) A) Simulation assuming zero-order synthesis and first-order degradation of ribosomes with an initial half-life of 7 days. Synthesis or degradation rate changed in simulation from baseline at t=0 by amount indicated; new and old ribosome levels represent all ribosomes present before or synthesized after a 9-day timepoint, to mimic the screen conditions. B) Highlighting genes included in sublibrary. All isoforms of a gene chosen for inclusion are included. Some genes with relatively high fold change but low counts were not included in the sublibrary construction. C) Score construction. Plots show individual sgRNAs; light grey shows non-targeting controls, dark gray are targeting sgRNAs, plotted by listed log_2_ fold change as listed. Lines and arrows show line fitted for score development. Scores are distance from zero parallel to line squared divided by sum of distance parallel to line and distance perpendicular to the line, providing a net score of distance parallel weighted by fraction of total distance from line that is parallel. Scores are shown by color as listed in colorbar. D) Pearson correlation of gene-level log_2_ fold change. Genome-wide libraries are subset to only include genes in sublibrary. E) GO categories enriched for positive or negative gene scores Adjusted p-value is based t-test upon Benjamini-Hochberg correction for t-test of univariate linear model coefficient for membership in GO categories (see methods, Table S6). GO category names are colored based upon categories listed at bottom right. F) Pearson correlation between synthesis and degradation scores and growth scores for log_2_ fold from sublibrary screens.

**Figure S3:**
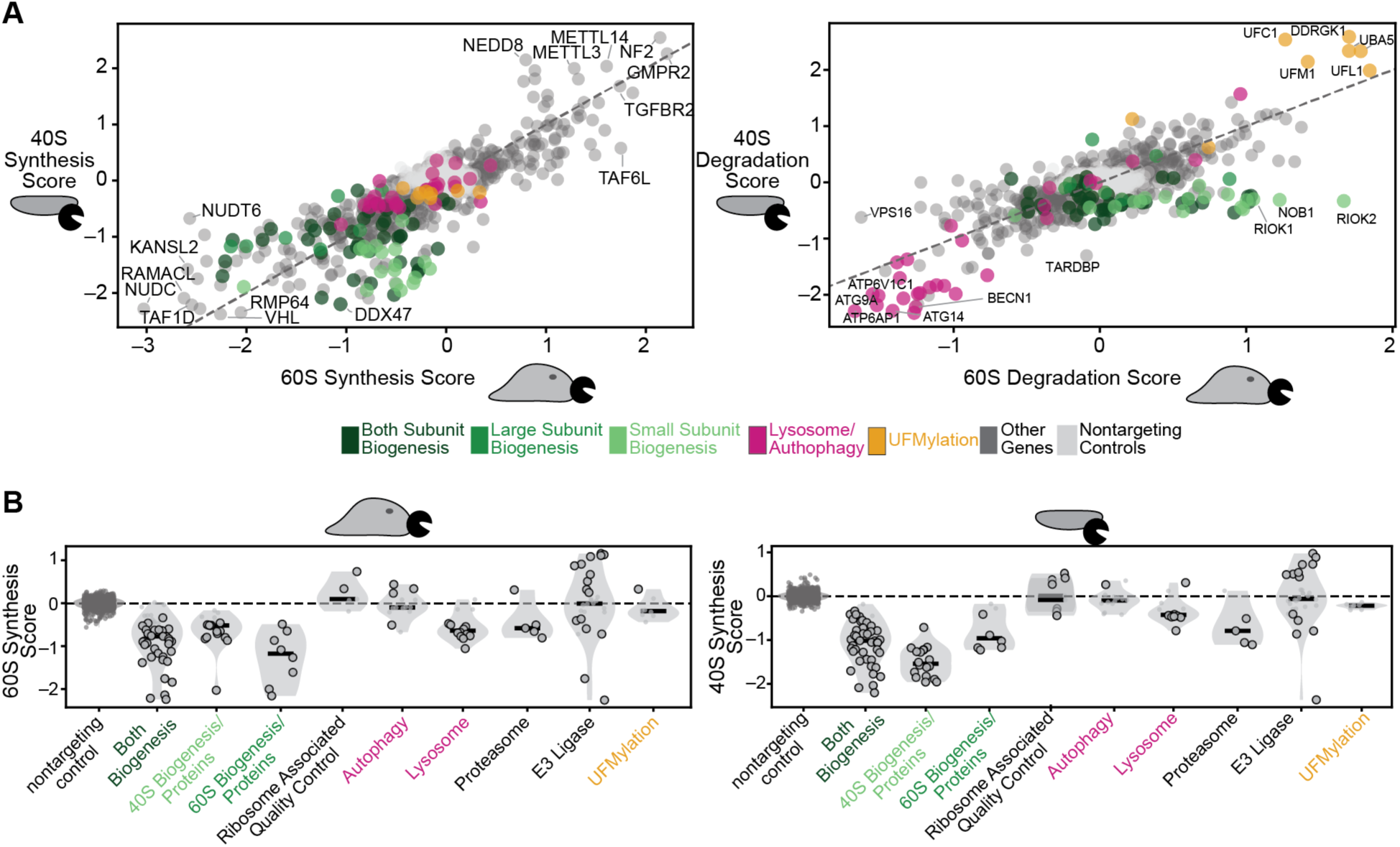
Gene and pathway level synthesis and degradation score effects (Related to Figure 2) A) Right: comparison of gene-level synthesis scores for 40S subunit (Y-axis) and 60S subunit (X-axis)-positive scores imply gene knockdown leads to faster ribosome synthesis. Left: Comparison of gene-level degradation scores for 40S subunit (Y-axis) and 60S subunit (X-axis)-positive scores imply gene knockdown leads to faster ribosome degradation B) Effects of specific gene categories on synthesis score for large subunit (left) and small subunit (right). Circled dots pass the 5% false discovery rate threshold compared to nontargeting controls by Mann-Whitney U-test.

**Figure S4:**
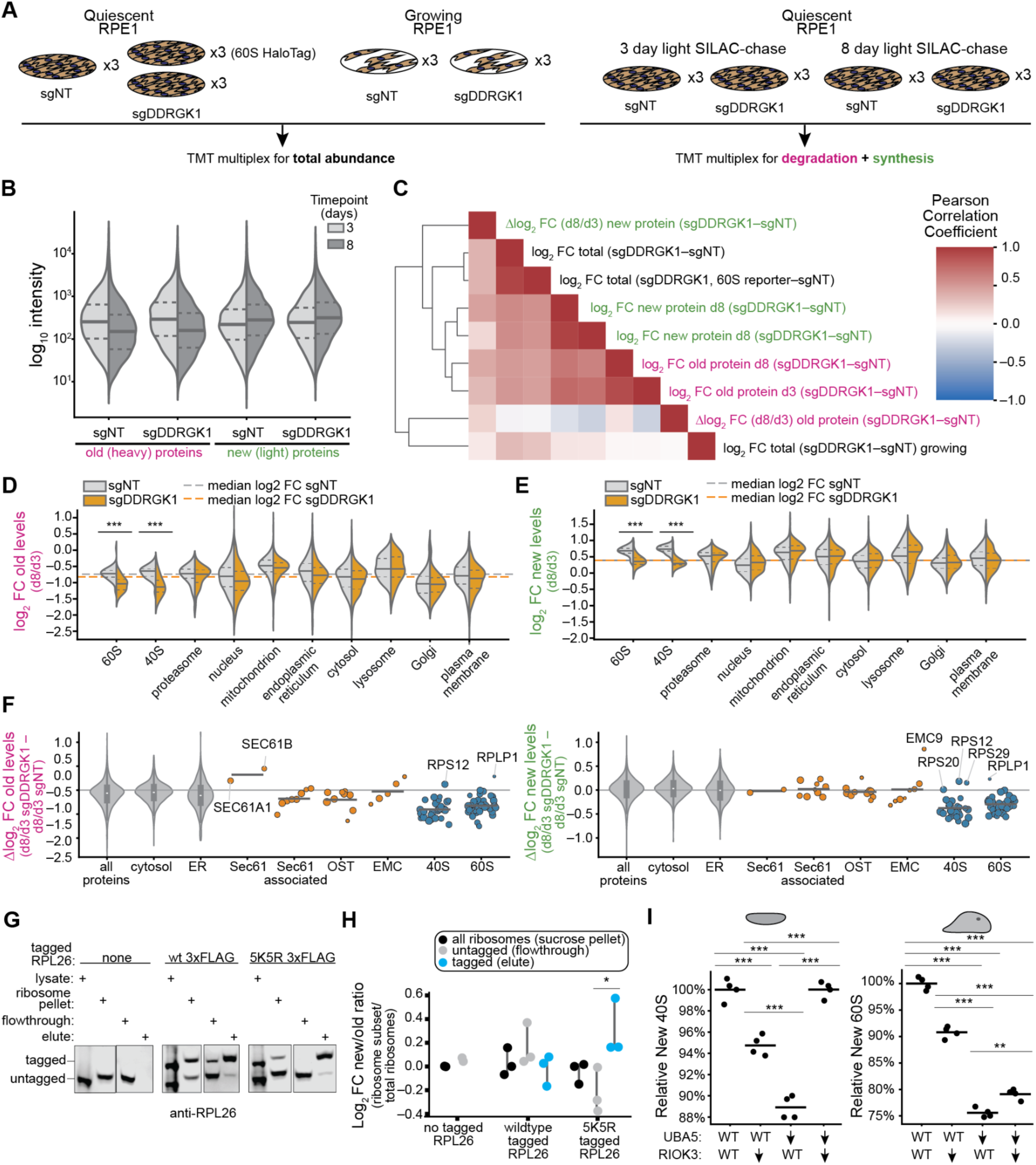
Proteome-level and targeted assays show accelerated ribosome degradation upon loss of UFMylation (Related to Figure 3) A) Schematic of proteomics datasets collected. Left: TMT multiplex of whole-cell proteomics (no SILAC labeling) across growing and quiescent cell states, as well as SILAC-chase samples. B) Changes in total protein intensity for old and new (heavy and light) proteins as measured by genotype C) Pearson correlation of total proteome log_2_ fold change between collected samples. Numbers represent an average of 3 replicates. Note separate TMT multiplex used for total protein abundance limit accuracy of direct comparisons to SILAC data. D) Change in old protein level split out by genotype, by cellular compartment. Relative change on specific proteins shown in Figure 3B. E) Change in new protein level split out by genotype, by cellular compartment. F) Change in new or old protein level of ER protein categories. Larger point size indicates higher MS signal. Horizontal grey lines are the average of given categories; violin plots are for categories with >100 members. G) RPL26 immunoblots showing representative images of different gel fractions. Note shifted bands due to salt content for lysate before sucrose pellet. Fractions during immunoprecipitation indicated above gel, as well as if over-expressed RPL26-3xFLAG is present and which variant. H) As in Figure 3F, but showing ribosome pellet in-gel fluorescence ratio before pulldown. All data normalized to the average of untagged ribosome level values. I) Change in new (since start of 5 day turnover experiment) ribosome level for 40S subunit (left) and 60S subunit (right) after RIOK3, UBA5, or both knockdown. **: p< 0.01, ***: p< 0.001, Welch’s t-test.

**Figure S5:**
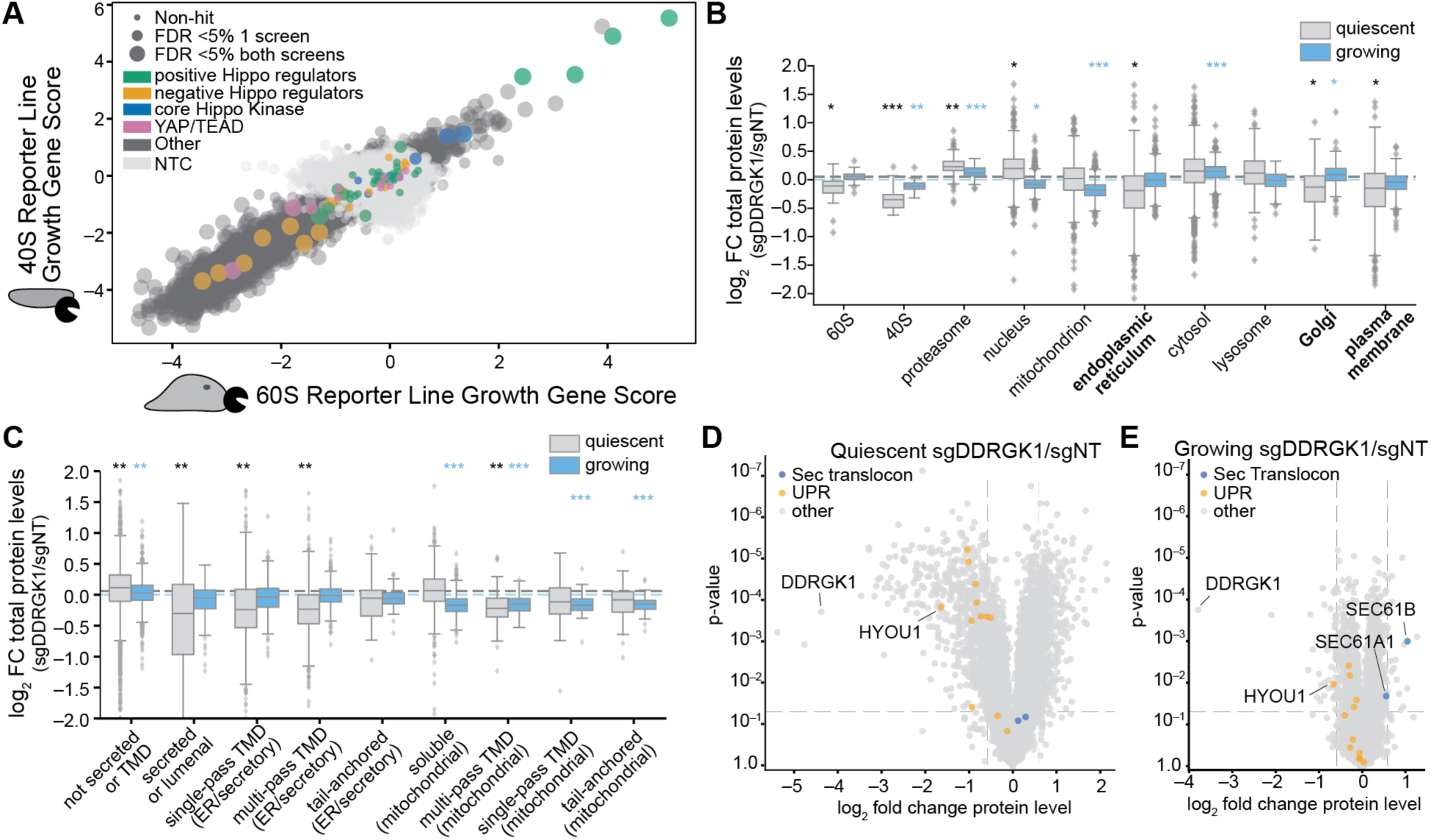
Contact-inhibited cells show greater defects in secretory pathway proteins upon UFMylation loss (Related to Figure 4) A) Comparison of genome-wide growth data from quiescent cells, highlighting Hippo pathway components known to respond to cell density, as well as core machinery (annotated in Supplementary Table S2). Gene score represents average log_2_ fold change in relative abundance between early and late timepoints, with positive scores representing increased guide abundance at late timepoints during quiescent cell screen. Point size represents significance testing passing the 5% FDR threshold in either or both screens. B) Change in protein levels upon long-term DDRGK1 knockdown for categories of proteins from quiescent and growing cells. Dashed lines show the median of all genes. *: p <0.05, **: p < 0.01, ***: p<0.001, Mann-Whitney U vs. rest of proteome, Benjamini-Hochberg corrected (categories displayed in Supplementary Table S9) C) As in panel A, but showing categories of membrane proteins. Categories displayed in Supplementary Table 9. D) Volcano plot showing average log_2_ fold change in protein level and p-value in t-test between DDRGK1 knockdown and control guide in quiescent state. ER-resident UPR-induced genes and components of Sec61 translocon highlighted; dashed lines show 1.5-fold change or p=0.05 level. Genes highlighted in Table S9 E) As in panel D, but showing comparison of knockdown in growing RPE1s at 10% serum.

**Figure S6:**
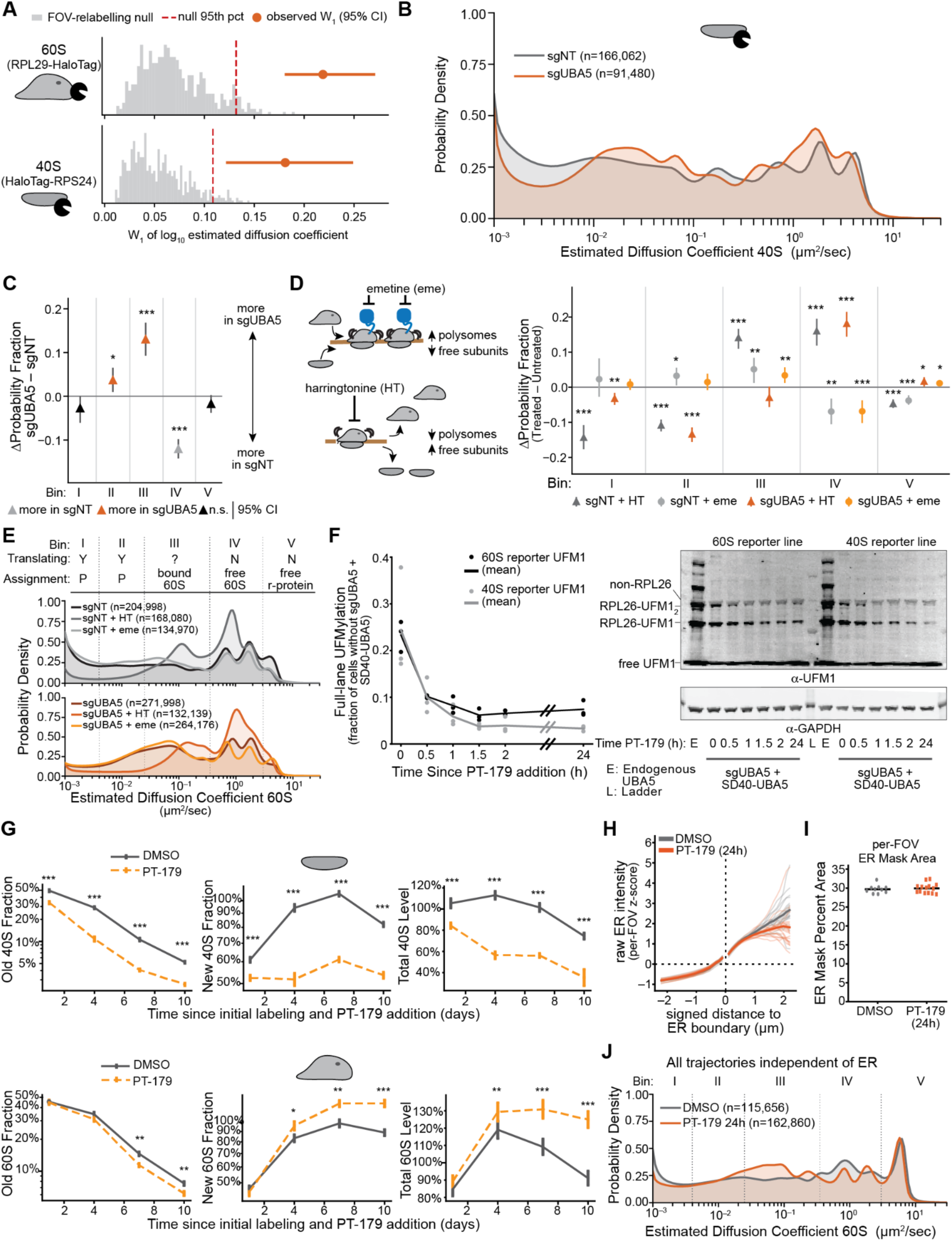
60S ribosomes accumulate in low-translation ER-bound state upon UFMylation loss (Related to Figure 5) A) Wasserstein distance (W1) between sgUBA5 and sgNT (10 FOV each) for large and small subunits. Range: 95% confidence interval based upon bootstrap resampling of FOVs and distribution recalculation. Red line-95% distance from random permutation of FOVs. B) Estimated diffusion coefficient from analysis of 40S HaloTag (Halo-RPS24). Colored by genetic knockdown; numbers represent number of subtrajectories analyzed for distributions C) Quantification of differences in diffusion bin integrated probability density for 60S ribosome from Figure 5C. Ranges represent 95% confidence intervals based upon bootstrap resampling of FOVs. Coloring shows if favors sgNT (grey) or sgUBA5 (orange)-bins not significant at p<0.05 using FOV permutation test. Grey horizontal line to show no difference between genetic conditions. D) Schematic and diffusion bin density for polysome run-off or trapping experiments in endogenously tagged 60S ribosomes. Harringtonine (10 ug/mL) or emetine (360 µM) was added for 1 hour prior to imaging. n= 20 FOV per condition. *: p < 0.05, **: p < 0.01, *** p < 0.001, Benjamini-Hochberg corrected from 2000 permutations of FOVs between conditions. Only carried out statistical comparison of drug treatment to untreated within-genotype or cross-genotype for same-treatment. E) Same data from panel D. Table at top shows conclusions in translation based upon increase or decrease upon drug treatment, and concluded species based on translation and literature diffusion rates (P: polysome, free: unincorporated protein; Y: translating, N: not translating; ?: ambiguous from data, initiation dependent sgUBA5, not sgNT). Bottom shows estimated 60S diffusion rate (x-axis) versus probability density from saSPT for sgUBA5 (bottom) and sgNT (top) treated with indicated drugs. F) Degradation efficiency of SD40-UBA5 construct expressed in sgUBA5 background. Data based on quantification of immunoblotting, n=3 replicates-representative blots shown on right. 20 µM PT-179 added at t=0. E: no UBA5 knockdown. All data collected after 10 minutes treatment with 10 nM Anisomycin to induce UFMylation upon ribosome collision. G) Changes based on flow cytometry in new, old, and total ribosome levels upon PT-179 treatment for indicated lengths. *: p < 0.05, **: p < 0.01, *** p < 0.001 Welch’s t-test comparison between DMSO and PT-179. Data normalized to average DMSO-treated total fluorescence. Ranges show standard error of the mean for n=4 replicates. H) Comparison of ER signal intensity based on distance to ER mask boundary in both conditions. Bold lines represent average across FOV (n= 10 DMSO, n=14 PT-179 treated), thin lines represent average of individual FOV. I) Total area per-FOV (and average) of ER mask under DMSO and PT-179 treatment. J) Total estimated diffusion distribution for all 60S ribosomes ignoring ER mask status (subset of data used in Figure 5F). In parentheses are the number of subtrajectories per condition.

**Figure S7:**
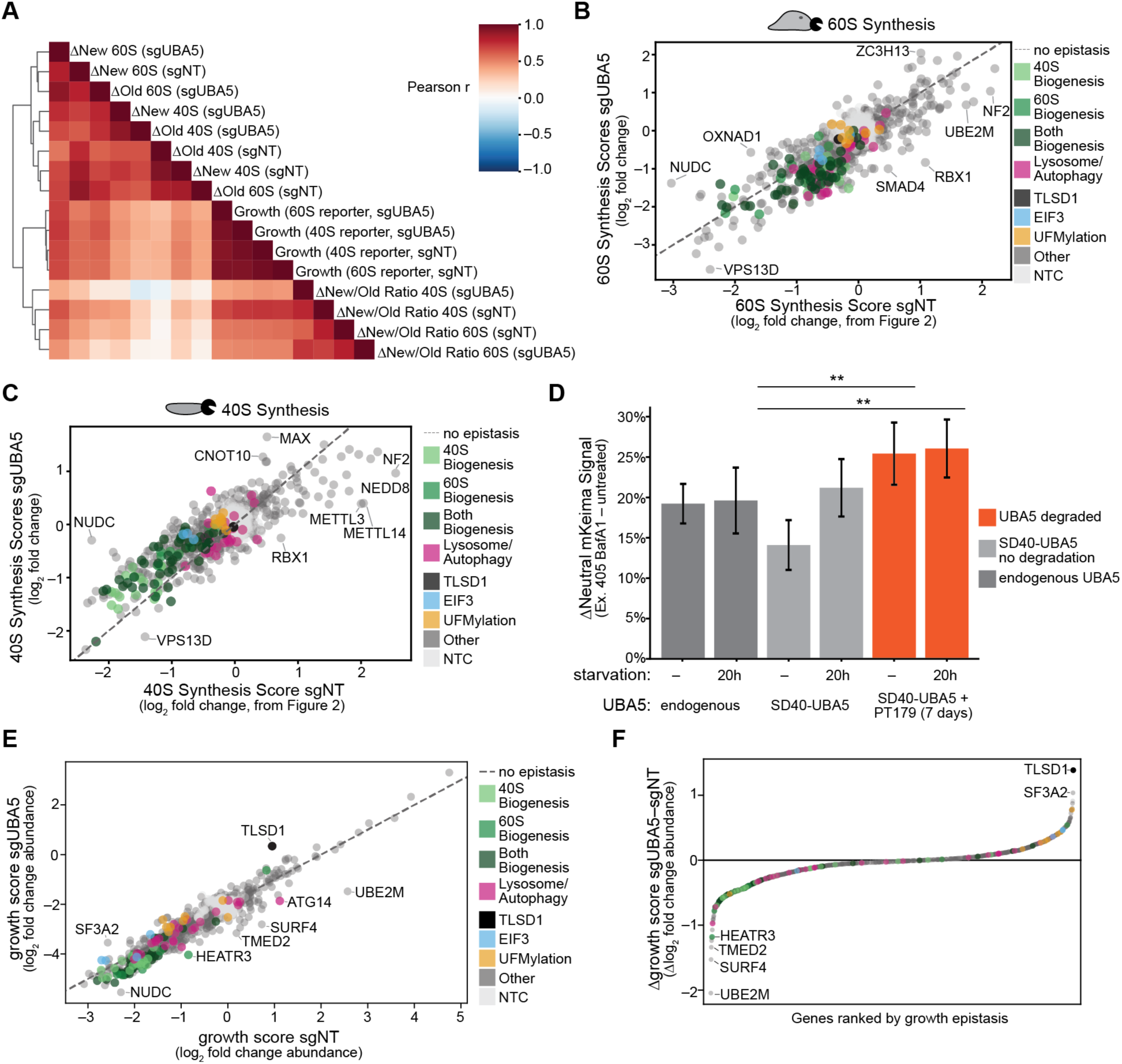
UBA5 knockdown shows limited epistatic effects on synthesis, growth, related to Figure 6 A) Correlation coefficient of sublibrary screens gene effects (log_2_ fold changes) with and without UBA5 knockdown B) Synthesis scores for 60S reporter for sgUBA5 screen (y-axis) and sgNT screen (x-axis), highlighting genes off-diagonal (dashed line shows expected additive effects). Colored by gene category, as in Figure 6B but including all genes. C) Synthesis scores for 40S reporter, as in panel B. D) Flow cytometry showing change in neutral signal from RPL29-mKeima upon overnight treatment with 10 nM Bafilomycin A1 (BafA1) under different conditions. Starved conditions were grown for 20 hours without serum or amino acids as a positive control for autophagy induction. **: p < 0.01, Benjamini-Hochberg corrected Welch’s t-test. Related to Figure 6C. n=6 samples per condition; bars are combined standard deviation of untreated and treated samples. E) Growth scores (averaged from RPL29-reporter and RPS3-reporter cell lines) for sgUBA5 and sgNT quiescent cell sublibrary screens. Genes showing off-diagonal growth effects highlighted, colored by gene category. F) Waterfall plot showing genes ranked by difference in growth score in sgUBA5 screen minus difference in growth score in sgNT screen. Genes ordered in x-axis by rank; difference shown on y-axis. Colored as in E, most extreme outlier genes labeled.

**Figure S8:**
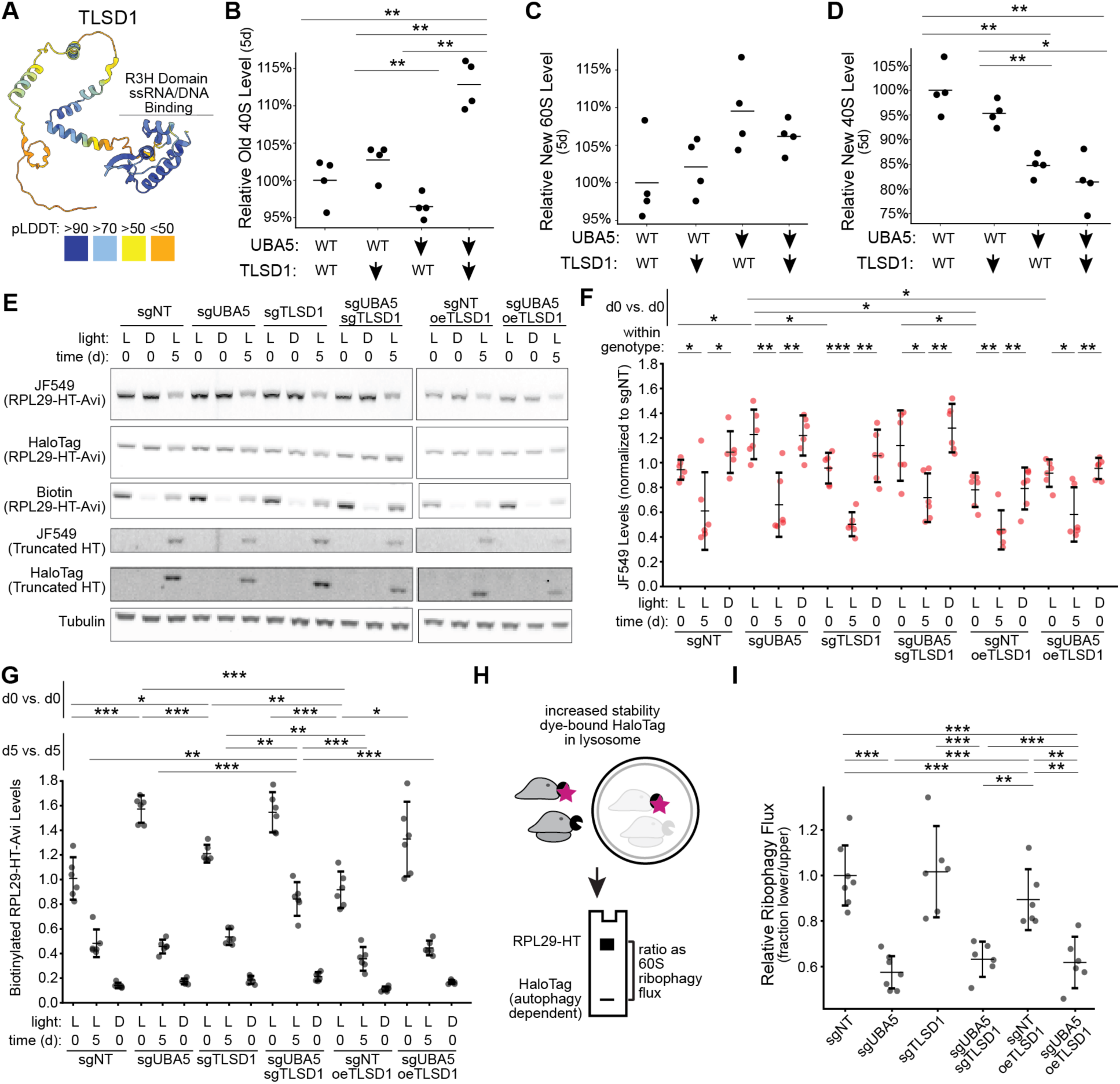
TLSD1 is induced by UFMylation loss and degrades ER ribosomes independent of autophagy, related to Figure 7 A) AlphaFold model of TLSD1, colored by pLDDT (metric of model certainty). R3H domain highlighted. B) Effect of UBA5 knockdown, TLSD1 knockdown, or both on >5-day old 40S ribosome levels by flow cytometry. Individual points represent median fluorescence intensity, normalized to mean of control (nontargeting guide). **: p < 0.01, Benjamini-Hochberg corrected from Welch’s t-test. C) Effect of UBA5 knockdown, TLSD1 knockdown, or both on <5 day old (“new”) 60S ribosome levels by flow cytometry. Individual points represent median fluorescence intensity, normalized to mean of control (nontargeting guide). **: p < 0.01, Benjamini-Hochberg corrected from Welch’s t-test. D) As in C, but for “new” 40S. E) Representative gel or immunoblot images for assay described in panels F–I and Figure 7B. F) Integrated JF549 signal from in-gel fluorescence, representing total initially-labeled RPL29-HaloTag from in-gel fluorescence, normalized to integrated tubulin level from fluorescent immunoblot, for samples containing RPL29-HaloTag-AviTag construct labeled with ER-targeted LOV-BirA followed by 1 hour labeling with 100 nM JF549-Halo dye. Lysate is collected either immediately after labeling (time (d) = 0) or 5 days later, as indicated below the graph. Light conditions either 15 minutes labeling on blue light with 500 nM biotin (L) or 500 nM biotin without blue light (D) to measure background signal in the biotin channel. All signals normalized to average sgNT 0d L signal; genetic condition indicated at bottom. oeTLSD1: TLSD1 overexpression. *: p < 0.05, **: p < 0.01 Benjamini-Hochberg adjusted p-values. Comparisons within-genotype and cross-genotype between 0d L conditions; no 5d L conditions met statistical significance. G) As in F, but showing integrated biotin signal for full-length RPL29-HaloTag-Avi (RPL29-HT-Avi) normalized to total tubulin signal as a loading control divided by JF549 signal from the same gel. Normalized to sgNT signal from zero day timepoint. Asterisks as above. Comparisons cross-genotype for same length and type of labeling; all within genotype comparisons met p<0.001 level of significance. H) Schematic of HaloTag lysosomal delivery assay. Lysosomal dye-labeled HaloTag is relatively stable, forming a lower-molecular weight band due to digestion of non-HaloTag portion of reporter prior to HaloTag itself. I) Quantification of ratio of lower band relative to upper band, normalized to mean of sgNT, as measure of 60S ribophagy. Only the 5 day timepoint is analyzed.

## Supplementary Tables

**Table S1: Genomewide Screen sgRNA-level data Table S2: Genomewide Screen gene-level data**

**Table S3: GO term enrichment from genome-wide screens**

**Table S4: sgRNA-level sublibrary data, including Synthesis and Degradation Scores Table S5: gene sublibrary data, including Synthesis and Degradation scores**

**S6: GO term enrichment for synthesis and degradation scores**

**S7: Targeted sgRNA spacer sequences and qPCR validation**

**S8: New/old SILAC TMT data, including compartment annotations**

**S9: total abundance TMT data, including multipass/secretory annotations**

**S10: sgUBA5 sublibrary screen data sgRNA level with scores**

**S11: sgUBA5 sublibrary screen gene-level data with scores**

**S12: Analysis parameters for single-particle tracking**

## Methods

### Cell culture and quiescence induction

hTERT-RPE1 (ATCC, CRL-4000, hereafter “RPE1”) were cultured in DMEM/F12 (Thermo Fisher Scientific, 11330032) supplemented with 10% fetal bovine serum (FBS; Avantor 97068-085) and 1:100 dilution of Penicillin/Streptomycin/Glutamine (PSG, Thermo Fisher Scientific, 10378016), at 37 °C and 5% CO₂. Cells were passaged using TrypLE (Thermo Fisher Scientific 12604013) every 2–5 days. Cells regularly tested negative for mycoplasma by qPCR. All derivative cell lines were generated from this base; only Figure 1 uses the base cell line.

For all quiescence experiments, cells were seeded at 3.5 x 10^5^ cells/cm^2^ and grown in appropriate vessels to full confluence a week, then switched to medium containing 1% FBS for a minimum of 5 days. For CRISPRi screens, Figure 1 turnover assays, and quiescent-state 20 µM S-trityl L-cysteine (STLC, Millipore Sigma, 164739-5G) in DMSO (40 mM stock) was added after these 5 days as a media component the night before HaloTag ligand addition, and continued to be added until the end of the assay. All other assays excluded this compound.

Assays carried out under growing conditions were maintained at 10% FBS and kept to less than 70% confluence during the duration of the assay, passaging as needed. Assays measuring translation directly in confluent cells (single-particle tracking or polysome analysis) were carried out on cells allowed to grow to full confluence but maintained at 10% serum for at least 5 days.

### Flow Cytometry and Fluorescence-activated Cell Sorting (FACS)

Flow cytometry data collection was performed on an Attune NxT flow cytometer (Thermo Fisher Scientific) equipped with autosampler, and analyzed using FlowJo 10.8.1 to obtain median fluorescence intensity from a minimum of 1000 singlet cells positive for BFP and GFP (CRISPRi machinery and sgRNA guide vectors respectively). All cell sorting for clonal populations carried out using a Sony MA900 cell sorter, sorting into 96-well plates containing complete medium. Polyclonal sorting carried out using either Sony MA900 or BD FACSAriaII in purity mode. Screen samples sorting is described below.

### Molecular Cloning and Construct Table

**Table 1:**
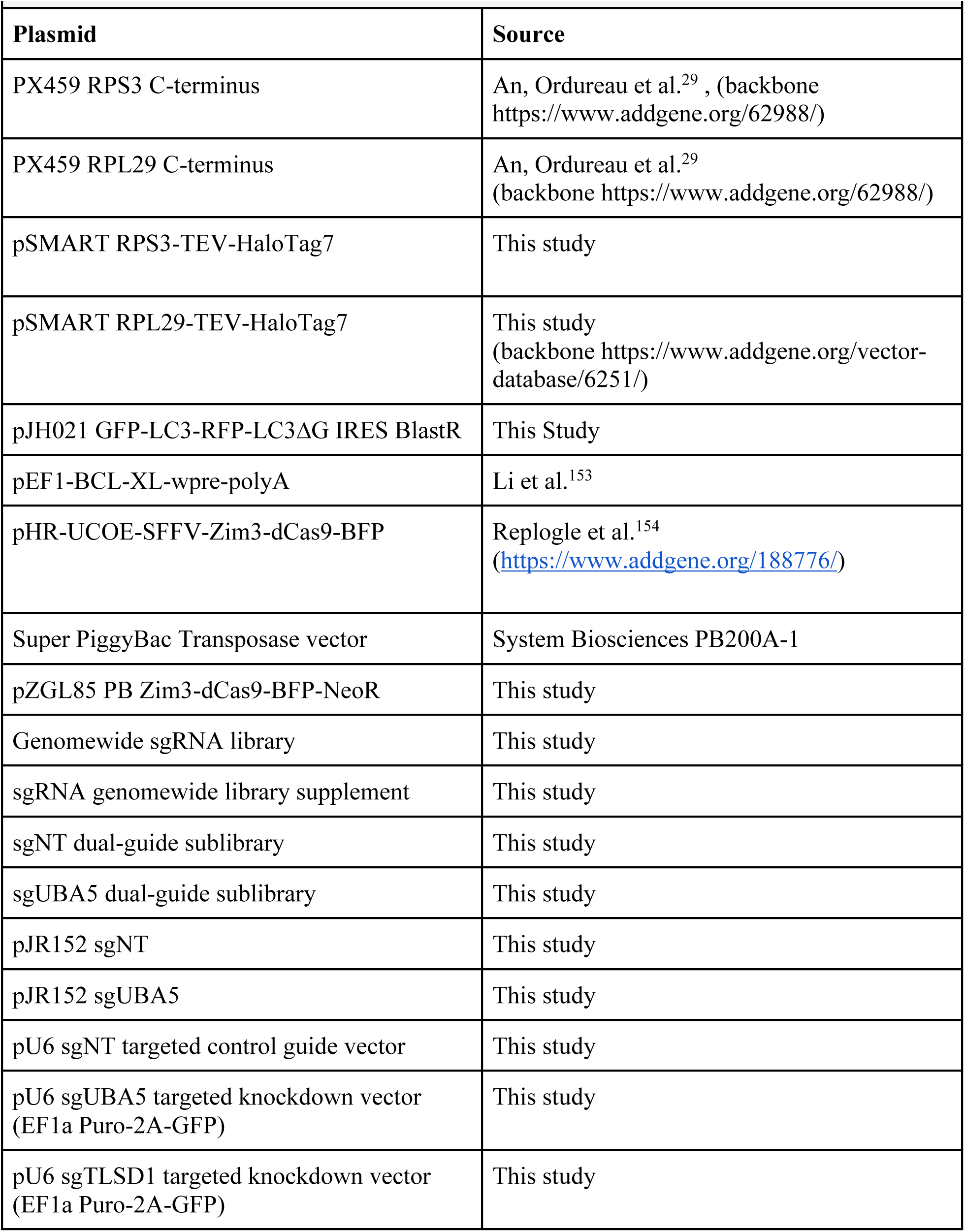
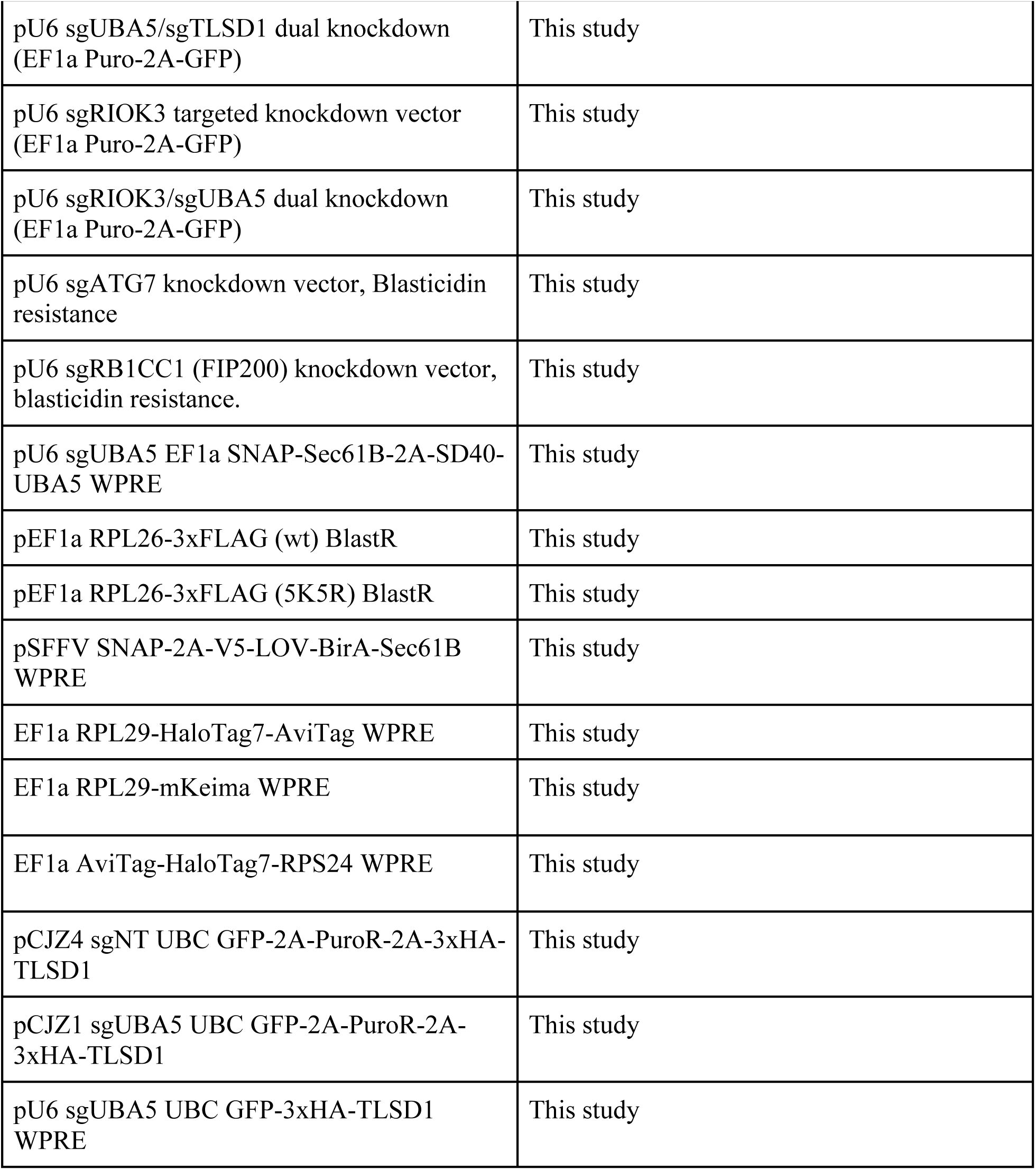
Plasmids used in study.

All targeted guide RNA plasmids were cloned from annealed oligos and inserted into a lentiviral pU6-sgRNA EF1a-Puro-T2A-GFP vector (Addgene 111596)^155^ digested with BlpI/BstXI. Dual guide vectors were cloned according to previously reported strategy.^113^ For autophagy knockdown vectors, Blasticidin resistance cassette was ordered as gBlock from IDT and used to replace Puro-2A-GFP via Gibson cloning. Protospacer sequences are reported in Supplementary Table 7 with single-guide knockdown validation. RPS3 and RPL29 HaloTag knock-in donor templates were modified to introduce TEV protease site into linker via Gibson cloning (New England Biolabs E2621). GFP-LC3-RFP-LC3ΔG autophagy reporter was subcloned from previously reported retroviral vector (kind gift of Dr. Noboru Mizushima, Addgene #84572) into pAG27 to generate lentiviral constructs used in this study via Gibson cloning. Neomycin resistance and Zim3-dCas9-mTagBFP2 were subcloned into piggybac vector (kind gift of Dr. Michael Ward, Addgene 204723). RPL26-3xFLAG mutant sequences were generated from Ensembl sequence RPL26-213 (transcript ENST00000648839), and 3xFLAG was appended to C-terminus with a GS linker; DNA for variants was synthesized as gBlocks by IDT and cloned along with IRES Blasticidin resistance into pCDH-EF1-IRES-GFP (kind gift of Dr. Oskar Laur, Addgene 128059) via Gibson cloning. SNAP-2A-V5 cassette was synthesized as a gBlock by IDT and subcloned into pJL95 (Addgene 243683) via Gibson cloning. AviTag-modified RiboHalo proteins were purchased as gBlocks from IDT (RPS24: Ensembl transcript ENST00000372360, RPL29: Ensembl transcript ENST00000294189), with M-AviTag-GGGGS-HaloTag7-GSGGS-TEV cleavage site-XTEN16 added in place of initiator methionine for RPS24 or XTEN16-TEV cleavage site-GSGGS-HaloTag7-GGGS-AviTag-stop codon in place of native stop codon. For RPL29-mKeima, the same construction was used but with XTEN16-mKeima (sequence derived from Addgene 131626)^156^ and cloned by Gibson cloning. GFP-3xHA-R3HDM4 (TLSD1) and GFP-2A-3xHA-R3HDM4 were cloned via Gibson cloning into pU6-sgRNA EF1a-Puro-T2A-GFP from IDT gBlocks using sequence from Ensembl transcript ENST00000361574. pJR152 vectors (Addgene 196280) were cloned as described previously.^113^

### Lentivirus infection and selection of RPE1s

All lentivirus was produced in HEK 293T C17 cells (ATCC CRL-11268) according to previously reported protocols.^157^ RPE1 cells were infected by plating into DMEM/F12 with 8 µg/mL polybrene (Millipore Sigma TR-1003-G) and appropriate concentrations of virus to achieve a multiplicity of infection (MOI) of approximately 0.3 at 35,000 cells/cm^2^; virus was removed after overnight incubation and cells maintained in DMEM/F12 with 10% serum and Pen/Strep/Glutamine for 48 hours before selection. All viruses were titered before use via fluorescence or antibiotic kill curve. Puromycin (Thermo Fisher Scientific A1113803) was used at 10 µg/mL for selection over 5 days with at least one passage, and Blasticidin S (Thermo Fisher Scientific A1113903) was used at 50 µg/mL for 10 days and 3 passages. Selection efficiency was confirmed and checked via flow cytometry.

### Library construction

Genome-wide sgRNA library was selected from our hCRISPRi v2 library^158^ using a three-tiered approach balancing empirical screen data with predicted rankings as described previously,^63^ yielding 61,581 sgRNAs targeting 20,527 transcription start sites (TSSs) of 18,903 genes (3 gRNAs/TSS) and 3,078 nontargeting controls. Genes from NCBI RefSeq Release 212 and GENCODE V41 annotations not included in the initial library (706 genes, 734 unique TSSs, 2202 guides) were added in a separate library to add coverage for these genes, choosing guides as described below using Dolcetto CRISPRi library design.^159^ Both parts were cloned into a plasmid derived from Addgene #187243^154^ by replacing the sgRNA scaffold with the F+E sgRNA scaffold as previously described.^158^

For sublibrary, top 715 genes were selected manually based upon effect size and differential effects between subunit. Some top-scoring genes were omitted due to lower total counts in genome-wide screen leading to exclusion from preliminary analysis; these genes were included upon re-analysis for completeness. For each gene, 3 further nonredundant guides per gene were pulled from the Dolcetto CRISPRi library design^159^ using updated NCBI RefSeq Release 212 (filtered for curated, experimentally validated transcripts) and GENCODE V41 annotations downloaded from the UCSC genome browser. New spacer candidates were filtered for not being in the genomewide library and containing BstXI or BlpI sites; 3 extra sites were selected per guide RNA, bringing number of guides to 6 per TSS; all TSSes for top scoring genes were used. 493 control sgRNAs were added to library, bringing total number of elements to 4928, ordered from Agilent and cloned into the same vector as the genomewide library. To make sublibrary compatible with dual-guide screening for epistasis, amplified library was digested with BamHI and NotI; we note that this removed 53 designed spacers, affecting 38 genes and 2 nontargeting controls. sgNT (wildtype screen, Figure 2) and sgUBA5 (suppressor screen, Figure 6) were inserted into library from pJR152 vectors as described in Guna et al.^113^

### SDS-PAGE and Immunoblot Protocols

Unless otherwise noted, cells growing in a 6-well plate were washed in ice-cold PBS and lysed with 1x RIPA buffer (Millipore Sigma R0278) containing Roche EDTA-free protease inhibitor, 100-200 µL. Lysate was clarified at 4°C at >4000 rcf, and normalized by Pierce BCA protein Assay (Thermo Fisher Scientific). Lysates were denatured using Laemmli buffer at 95°C for 5 minutes, and loaded in equal amounts onto 4-12% Bolt Bis-Tris gels (Thermo Fisher Scientific NW04125BOX) or 12% Bolt Bis-Tris Gels (Thermo Fisher Scientific NW00127BOX) as described per-assay, separated via electrophoresis at 150V in MOPS buffer (Thermo Fisher Scientific B000102). In gel fluorescence, if performed, was carried out on a Typhoon Fluorescence FLA9500 scanner (GE Healthcare). For immunoblotting, proteins were transferred onto 0.2 µm nitrocellulose (BioRad 1704270) or 0.42 µm PVDF (BioRad 1704274) membranes (activated using methanol) using TransBlot Turbo (BioRad) on mixed molecular weight settings according to manufacturer’s instructions. Membranes were blocked for one hour at room temperature using Intercept TBS blocking buffer (LICORbio 927-60001), incubated overnight with primary antibodies (listed below) diluted 1:1000 in SuperBlock (TBS) T20 (Thermo Fisher Scientific 37536). After washing at room temperature with TBS with 0.05% Tween-20 (TBS-T), LICORbio secondary antibodies (IRDye 800CW goat anti-mouse, 926-32210, goat anti-rabbit 926-32211, IRDye 680LT Donkey anti-rabbit 926-68023, and IRDye 680LT goat anti-mouse 926-68020) were used at 1:10,000 dilution in SuperBlock T20 with 0.01% SDS for 1 hour, followed by washing with TBS-T and imaging on a LICOR Odyssey CLX scanner. RPL29 was imaged using anti-mouse HRP (BioRad 1721011) at 1:10,000 dilution in Superblock T20 and imaged with an iBrightCL750 imager (Thermo Fisher Scientific A44116).

Primary antibodies: RPS3: Rabbit monoclonal D50G7 (Cell Signaling 9538), RPL29 Abnova MaxPab polyclonal H00006159-B01P, HaloTag Promega Mouse monoclonal G9211, UFM1 Abcam rabbit monoclonal EPR4264(2) (ab109305), RPL26 abcam rabbit polyclonal ab59567, beta-Tubulin rabbit monoclonal Cell Signaling Technologies (2128), GAPDH mouse monoclonal D4C6R Cell Signaling Technologies (97166S), Steptavidin IRDye680LT LICOR Biosciences (926-68031).

### qPCR Validation of sgRNAs

Performed as previously described in detail^160^ in confluent RPE1s. sgNT knockdown cells used as control. qPCR probes used listed in Supplementary Table S7. qPCR performed using QuantStudio 7 Flex (Thermo Fisher Scientific).

### Endogenous HaloTag Editing and Screening Cell Line Construction

RPE1 RiboHalo screening cell lines were constructed via HDR. RPE1 were plated at 100,000 cells per 6 well dish 1 day prior to transfection in DMEM/F12 with 10% FBS without antibiotics. DNA mixture of PX459 containing guides targeting RPS3 (protospacer 5′-GACATACCTGTTATGCTGTG-3′) or RPL29 (protospace 5′-GAGATATCTCTGCCAACATG-3′) (700 ng) and corresponding HDR donor plasmids containing a C-terminal HaloTag fusion (700 ng) alongside BCL-XL-expressing plasmid to improve survival (100 ng) were mixed in 200 µL OptiMEM (Thermo Fisher Scientific 31985062) with 4 µL Lipofectamine Stem (Thermo Fisher Scientific STEM00008), incubated 10 minutes at room temperature, and added dropwise to cells. Medium was changed 24 hours later, and after 4 days cells were stained with JF646-HaloTag dye and single cell isolated by FACS into 96 well plates; clones were expanded, tested for fluorescence, and genome editing was confirmed by immunoblot on 4-12% gels from whole cell lysate (RPS3) and ribosome pellet (RPL29, due to off-target bands). To introduce CRISPRi machinery, lentivirus containing SFFV Zim3-dCas9-2xNLS-BFP infected these cells, and clones were isolated via single-cell sorting.

CRISPRi knockdown efficacy of clones confirmed via surface staining paired with knockdown of cell surface markers before use in screening and targeted assays. Cell lines described here and derivatives thereof used in all experiments in Figures 1–4, Figure 6B, Supplementary Figures 1– 5, Figure 7A and C, Supplementary Figure 6D–E, Supplementary Figure 7A–C and E–F, and Supplementary Figure 8B–D. Other cell-based assays used the polyclonal cell lines reported below to ensure effects were not due to clonal isolation artifacts.

### Polyclonal RiboHalo-AviTag and RPL29-mKeima Cell Line Construction

RPE1 cells were co-transfected with pZGL85 PB Zim3-dCas9-BFP-NeoR and Super PiggyBac transposase at a 10:1 ratio with 2 µg added in 6-well using transfection conditions described above for genome editing. After 72 hours, cells were FACS sorted for >100,000 cells with high BFP expression, then selected with Geneticin (Thermo Fisher Scientific 10131035) at 0.5 mg/mL for 10 days and 3 passages. Lentivirus encoding RPL29-HaloTag7-AviTag, AviTag-HaloTag7-RPS24, or RPL29-mKeima were used to infect these polyclonal RPE1 cells and then the infected cells were sorted (with JF646 staining for RiboHalo constructs) for expression and expanded. Frozen stocks were further thawed and infected with guide RNAs before use.

### EdU labeling assay

EdU labeling was carried out using Click-IT Plus EdU Flow Cytometry Assay Kit (Thermo Fisher Scientific C10634) according to manufacturer instructions with 1 µM EdU in complete media for growth conditions for 24 hours before collection and staining. Cells were dissociated using TrypLE after PBS wash then fixed, permeabilized, and stained according to manufacturer protocol.

### HaloTag pulse-chase turnover assay by flow cytometry

All HaloTag dyes were used at 100 nM, an amount that we found to saturate the signal and prevent subsequent staining with a counter dye in cells or after lysis (data not shown). Staining was performed at 37°C and 5% CO_2_ in a humidified incubator.

For initial testing and obtaining of turnover curves presented in Figure 1D, 18 wells of a 96-well plate were plated per timepoint on each RiboHalo endogenously edited cell line before introduction of CRISPRi machinery, with a separate plate for each timepoint, and unused wells filled with 200 µL PBS to reduce edge effects. Cells were grown to confluence and serum reduced to 1%. After 4 days at 1% serum 20 µM STLC was added to media and maintained for the entire experiment. Under each condition, 9 wells per timepoint were stained with JF549-Halo or JF646-Halo (18 wells total per timepoint) for 1 hour. Cells were then washed three times with media, media changed after a 1 hour incubation to remove residual dye that was intracellular, and incubated for the indicated amount of time until the endpoint. At the endpoint, 3 wells stained with the opposite dye from their initial staining, and 6 stained with the same dye as initially as fully-labeled controls, adding 100 nM dye for 1 hour, followed by 3 washes and 1 hour incubation in dye-free media. Cells were then washed with PBS, dissociated with TrypLE, then cell triturated, transferred to a fresh round-bottom 96 well plate, and acquired via Attune, with JF549 measured in channel “YL1-A” and JF646 in channel “RL1-A”. Data was aggregated as median fluorescence intensity (MFI) of ≥500 cells per well after gating for single viable cells using forward scatter and side scatter. Only cells positive for HaloTag staining were analyzed, providing a further removal of cells that had undergone membrane permeabilization. MFI from dual stained wells was divided by average of same-day singly stained controls for each dye, producing *F_remainin_*_g,*preliminary*_ and *F_ne_*_w,*preliminary*_ for each well. We calculated *F_remainin_*_g_ = 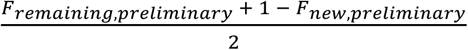 for each point. Standard error of the mean (sem for n=6) from controls and multiply stained wells was combined as standard error of the mean via root-sum-square method. We note that the lack of immediate chase likely led to some residual labeling at early timepoints, leading to fits >1 when extrapolating to t=0; we shifted to chasing with the second dye at the time of washout for subsequent experiments.

For subsequent targeted turnover experiments (Figures 3H, 4B–D, 7A, S4I, S6G, S8B–D), experiments were carried out in 24-well plates, except low-confluence experiments presented in figure 4, which were carried out in 6-well plates to boost cell numbers. Staining and chase were performed as described above, but consistently using JF549-Halo as the first stain, with JF646 as the chase. In addition to gating on singlets, all cells gated on GFP and BFP to ensure sgRNA and dCas9 expression. Single-staining controls were only carried out for the experiments in Supplementary Figure 6G (testing effects of SD40-UBA5 degron); for all others, new (JF549) and old (JF646) levels were directly analyzed relative to other genotypes at the same timepoint, and normalized to average MFI from sgNT controls. Degron characterization normalized to average singly-stained DMSO control across all timepoints for each dye to normalize, and error from this average propagated as root sum squared of standard error of mean from background and replicates per timepoint. Total for each sample at timepoint is n=12 points, including 4 singly-stained controls for each dye and pulse-chase samples, reported as sum of dyes normalized to single-stained DMSO-control averages, which centers on a 100%. GNE-7883 (Selleck Chemicals E1563), when added, was added at start of turnover experiment All statistical comparisons carried out as Welch’s t-test with Benjamini-Hochberg correction for multiple comparisons across genotypes in a single experiment.

### CRISPRi screen experimental protocol

At every split described below, all cells were pooled then re-distributed into plates, to maintain library diversity per-plate. Dye addition was performed by adding dye to relevant media and incubating for one hour in a humidified incubator at 37°C and 5% CO_2_.

Lentivirus was produced as described above in sufficient quantity to infect cells at coverage (Genomewide: 1000x coverage (63.7 million cells per replicate, two replicates x two cell lines) Sublibrary: 2000x coverage (9.75 million cells per replicate, two cell lines, two replicates each, with coverage for growth, new/old ratio, new, and old, giving 8000x coverage (39 million cells) per replicate per subunit)). Cells infected at MOI of 0.3 by combining with virus in media with 10% serum and penicillin/streptomycin/glutamine with 8 µg/mL polybrene, then plating at a density of 5 millions cells per 15 cm plate. At 24 hours, media was replaced with media without polybrene. At 48 hours after plating, 10 µg/mL puromycin was added, media changed to refresh puromycin daily. After 3 days of selection, cells were split, GFP positivity was measured at ∼85%, and 5 million GFP+ cells were plated per 15 cm plate, maintaining 2500x coverage and keeping 10 µg/mL puromycin. At this point, a cell sample at 1000x coverage was fixed with 4% paraformaldehyde for 45 minutes at 4°C, then frozen in 10%DMSO in FBS to provide an early timepoint for growth analysis.

For quiescent screens, media with puromycin was added for one final day. For genome-wide screens, one fifth of the plates below were used in the growing cell experiment the cells were maintained with media changes every 2-3 days until grown to confluence (at 10 days post-splitting, 14 days since initial infection). Cells then shifted to media with 1% serum and PSG and maintained in this media for four days, changing media after two days, then 20 µM STLC was added on day 5 of 1% FBS. The following morning, all cells were labeled by adding 15 mL 1% FBS media with 20 µM STLC and 100 nM JF549 Cells were washed in media without dye, then left in 20 mL media with 50 nM JF46. Media was changed thereafter every two to three days, then on day 9 after initial labeling 100 nM JF646 was added for 1 hour, washed with media, cells washed with PBS, then dissociated with TrypLE and filtered through 40 µm cell strainer. At this stage each plate was maintained separately through sorting. TrypLE was quenched with 5% FBS in PBS with 1 mM EDTA, cells were pelleted, resuspended in PBS, re-pelleted, then fixed in 4% paraformaldehyde in PBS at 4°C for 30 minutes. They were diluted 1:2 in FACS buffer (1% BSA in PBS with 1 mM EDTA), pelleted, frozen in FBS with 10% DMSO for sorting.

For growing arm of screen, at initial split, cells from same replicates were split into plates at 5 million cells per plate (an additional 1000x coverage total, 500x for growth and 500x for new/old ratio), then allowed to finish selection and grown for 5 days further before splitting again and replating at the same density. Cells were labeled with JF549 (100 nM) for 1 hour in media containing 10% FBS, followed by washing with media and adding JF646 (50 nM as chase dye). After 3 days, JF646 was re-added for 1 hour, cells washed and incubated for an hour without dye, then dissociated in PBS, filtered, fixed, and frozen separately for each plate as described for quiescent cell screens. Growing screens were performed as a single replicate split from the initial replicate of the genome-wide screen.

For genome-wide screens, frozen cells were split in half, with one half held in reserve as a late timepoint for growth. The remaining half (>1000x coverage of quiescent library, 500x for growing cell screen) was thawed, washed and resuspended in FACS buffer, then sorted per-plate on a BD FACSAriaII in yield mode for the top and bottom 25% each ratio of JF646/JF549 signal, gating on single cells based on forward and side scatter and requiring both GFP and BFP positivity. Cells from the same condition and sorting gate were pooled after sorting, keeping replicates separate, pelleted, and frozen for DNA extraction. For sublibrary screens a similar procedure was performed, but only one quarter of the cells (2 plates per screen) were reserved for growth analysis. The remaining three quarters were sorted per-plate on BD FACSAria II on yield mode, with two plates each sorted on top and bottom 25% of JF549/JF646, JF549 alone, and JF646 alone. Post-sorting cells were frozen as described for genome-wide.

For genomic DNA extraction, fixed cell pellets (post-sorting or frozen) were thawed and resuspended in lysis buffer (1% SDS and 10 mM EDTA in 50 mM Tris at pH 8.0) at a concentration of 20 million cells per mL. Mixture was aliquoted into 96 well plates in a thermocycler (100 µL per well), uncrosslinked at 65°C for 10 minutes, 2 µL Ambion RNAse cocktail was added per well (Thermo Fisher Scientific AM2286) and incubated at 37°C for 30 minutes, and proteinase K (10 µL per well) was added followed by 2 hours incubation at 37°C. Samples were cooled, re-pooled, and DNA purified using nucleospin blood (XL for genome-wide (Machery Nagel 740950) L for sublibraries (Machery Nagel 740954)) according to manufacturers instructions for cultured cells, with PBS used to bring samples up to appropriate volume for repeat proteinase K digest. After purification, DNA concentration was measured by qubit dsDNA high sensitivity kit (Thermo Fisher Scientific Q32851), and PCR reactions were prepared at 2.5 µg of genomic DNA per 100 µL, with 0.5 µM primer, using primers 3’-TCGTCGGCAGCGTCAGATGTGTATAAGAGACAG N_0–5_ CCCTTGGAGAACCACCTTGTTGG-5’ as forward and either 3’– GTCTCGTGGGCTCGGAGATGTGTATAAGAGACAG N_0–5_ CGGCCGCCTAATCCTGCA– 5’ (genome-wide library) or 3’–GTCTCGTGGGCTCGGAGATGTGTATAAGAGACAG N_0–5_ GGGAAATAGGCCCTCTTCCTGC–5’ (dual-guide sublibrary) as reverse, ordered from IDT, with N_0–5_ representing random nucleotide insertion of length 0–5 pooled to introduce diversity during sequencing. DNA was amplified with Q5 Ultra II Master Mix (New England Biolabs M0544) under the following conditions: 98°C for 3 minutes initial denaturation, 22 cycles of 10 seconds 98°C denaturation, 10 seconds 60°C annealing, and 25 seconds of 72°C extension, followed by a final 2 minute 72°C extension. PCRs were re-pooled and mixed per-sample, an aliquot was gel purified, and each sample was indexed with Nextera N700 and N500 series primers (5’–CAAGCAGAAGACGGCATACGAGATNNNNNNNNGTCTCGTGGGCTCGG–3’ and 5’– AATGATACGGCGACCACCGAGATCTACACNNNNNNNNTCGTCGGCAGCGTC–3’) (Illumina FC-131-2001) for 6 cycles. Samples were gel purified, quantified by qubit, and pooled at an equimolar ratio, then concentration confirmed by KAPA library quantification kit (Roche 07960140001) using QuantStudio #### qPCR machine. Libraries were sequenced using a NextSeq 2000 P3 100 cycles (genome-wide, Illumina 20040559) or XLEAP-SBS P4 (sublibrary: Illumina 20100995) with 5% PhiX spike-in (Illumina FC-110-3001) according to manufacturer’s protocols.

### CRISPRi Screen analysis

Sequencing data were de-multiplexed using bcl-convert and spacers counted using a custom python script, based upon exact sequence matches to spacer sequence library from pool exactly after the key sequence of CACCTTGTTG. For new/old ratio, new, old, and growth screens, counts were analyzed using the MAGeCK-iNC pipeline developed by the Kampmann laboratory (https://kampmannlab.ucsf.edu/mageck-inc).^66,67^ All 3 guides were taken into account for genome-wide analysis, while top 3 guides (based upon p-value for difference from MAGeCK) were used for sublibrary analysis, with a count threshold of at least 50 counts from combined treatment and control groups (top and bottom parts of sorting or early and late) for both replicates. All guide sequences in the library are present in Supplementary Tables S1, S4, and S10, with NaN for downstream analysis of guides not passing the threshold. For single replicate growth data, MAGeCK fits the mean-variance model to pooled treatment and control samples, inflating variance for these samples. Gene-level scores epsilon are mean log_2_ fold change in normalized counts, and p-values are Mann-Whitney U test comparing the two-sided p-values from MAGeCK for the sgRNAs to all non-targeting sgRNAs. Non-targeting control “genes” (NTC) are pseudogenes constructed by taking the 3 most-extreme-valued nontargeting controls from a randomly selected set of 5; process was repeated to construct the same number of NTC genes as true gene symbols. Mean epsilon of NTC was subtracted from all epsilon values to center data, and hits were set based on choosing the most extreme product of –log_10_ of p-value multiplied by epsilon that has ≤5% of “genes” passing threshold as NTC pseudogenes.

### GO term and pathway analysis

Gene-level scores were tested for GO-term enrichment by univariate linear modeling^161^ as implemented in the decoupler package using the 2025 Molecular Function, Biological Process, and Cellular Component GO term annotations^162,163^ as hosted by Enrichr.^164–166^ Linear modeling proved less biased towards large gene sets than other approaches; linear response was fit to epsilon for a given gene-level score.

Pathways were derived from the Proteostasis Consortium annotation with manual curation,^167–169^ and are listed in gene-level tables. Genes with bifunctional endogenous promoters with strong phenotypes that correspond cleanly to an enriched category are labeled for their bifunctional gene, as CRISPRi would inhibit transcription of both guides;^154^ these are denoted in gene names in tables, and include re-annotating ACAD11 as UBA5 and ANKZF1 as ATG9A in plots (both denoted in tables), as both have anomalous large phenotypes that perfectly correspond to their bifunctional genes.

### GFP-LC3-RFP-LC3ΔG bulk autophagy reporter

RPL29 and RPS3 screening cell lines were infected with guide RNA vectors against RB1CC1 or ATG7 and blasticidin selected for 20 days; screening cells also infected with reporter vector and blasticidin selected, with GFP positivity used to monitor selection efficacy. The autophagy-knockdown cells were infected with GFP-LC3-RFP-LC3ΔG lentivirus, and all reporter cells sorted for GFP and RFP positivity to remove lentiviral recombinants with a single fluorophore. For assay, cells were plated in 24 well plates (quiescent) or 6 well plates (growing or overnight starvation). Quiescent cells were grown to confluence and serum reduced to 1% for 5 days before assay; 10% FBS cells were grown at moderate confluence, and 16 hour EBSS cells were taken from this growing condition and subjected to 16 hour overnight growth in EBSS (Thermo Fisher Scientific 24010043) for amino acid starvation as a control for autophagy induction. Data were collected by flow cytometry after trypsin dissociation, and ratio of signal for GFP to RFP was calculated on a per-cell basis (after gating for single cells), with the median from the population for each well used for further analysis. For each growth condition, mean GFP/RFP ratio of control wells (n=3) was subtracted from mean GFP/RFP ratio of autophagy knockdown cells in the same growth condition. The difference plus standard error is plotted, and the differences are compared using Welch’s t-test.

### Turnover Simulation

We simulated biomolecule turnover using simple parameters, assuming zero-order synthesis and first-order degradation, *dN*/*dt* = *k_synt_*_ℎ*esis*_ − *k_de_*_g*radation*_*N*, where *k_synt_*_ℎ*esis*_ is synthesis rate, *k_de_*_g*radation*_ is degradation rate, and *N* is the number of biomolecules. Steady state abundance is *N_steady state_* = *k_synt_*_ℎ*esis*_ / *k_de_*_g*radation*_ and half life is *t*_1/2_ = *ln* (2)/*k_de_*_g*radation*_. Initial half life was set to 7 days, giving *k_de_*_g*radation*_ = 0.099 day^-1^molecule^-1^ with a synthesis rate of 100 per day. Molecules were subdivided as old or new if they were newer or older than this half life. In order to simulate this division, cohorts were simulated using a time step of 0.05 days, adding 5 new molecules and reducing cohort size by multiplying each by *e*^-*k*^*^degradation^*^∗0.05^. Fractional molecules were allowed, since simulating high-abundance ribosomal proteins; we acknowledge this simplistic simulation does not account for multiple exponential decay observed for complexes,^59^ but will accurately approximate the longer timescale degradation under study here.

Simulations modeled using Python, with numpy, matplotlib, and scipy packages. Code available upon request and will be deposited with publication.

### Synthesis and Degradation Score Analysis

Synthesis and degradation scores are calculated per-sgRNA based upon the log_2_ fold change calculated from the initial MAGeCK analysis, based upon a weighted metric using two perpendicular lines corresponding to *y* = *x* for synthesis and *y* =– *x* for degradation, where *y* is the log_2_ fold change in guide abundance between the top and bottom based on new/old ratio, and *x* is log_2_ fold change in guide abundance based either on new ribosomes (for synthesis) or on old ribosomes (for degradation). The final metric for each is *score* = *distance_parallel_* × 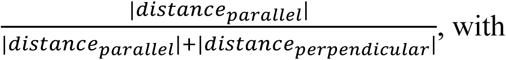

for synthesis:

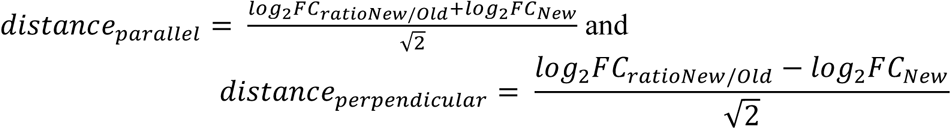

for degradation:

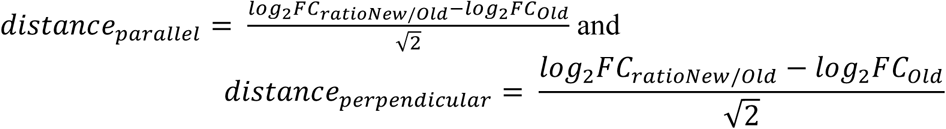

Scores were calculated for every sgRNA for each unless failing count thresholding as described below.

To assign per-sgRNA p-values, counts from all 8 samples contributing to the score (high and low sort gates on ratio and abundance, with 2 replicates), counts were median-of-ratios normalized, and guides with mean normalized count <20 were discarded (121-171 guides removed). A pseudocount of 32 was added and data were log_2_ transformed. Each guide’s per-population mean was subtracted to give residuals, which were pooled across all 4 populations to give a variance estimate with 4 degrees of freedom. To improve variance estimates and pool noise estimation, log_2_ variance was regressed against log_2_ of the mean normalized count with pseudocount of 32 added, and the obtained line of best fit was used to calculate an adjusted variance per-guide.This variance was used to calculate test statistic 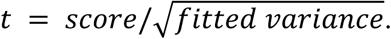 No distributional assumptions were made about these values-instead, the distribution of non-targetting guides was used as the null distribution to construct an empirical 2-sided p-value based on |*t*|, using the formula 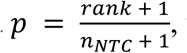 where rank is rank of guide with NTCs, 1 being. Score and p-value are reported in Supplementary Tables S4 and S10.

To generate gene-level scores, the same Mann-Whitney U protocol and synthetic null construction used from the MAGeCK-iNC pipeline was used with modifications. Top guides per gene were chosen based on minimum p-value, and ties were broken based on score, as top-scoring genes sat fully outside of empirical null distribution. Epsilon was the average of top 3 scores, and p-value was based on Mann-Whitney U rank test from test sgRNAs for a gene versus control sgRNAs. Non-targeting controls were selected per-gene based on drawing the a matched number of random non-targeting controls to guides for that gene (varies from 1–19 based on isoforms). Product is the product of epsilon and –log_10_ of p-value. This was carried out identically between sgNT and sgUBA5 screens. Epistasis between the sgUBA5 and control arms is reported in Supplementary Table S11 as the difference of gene scores (sgUBA5 − sgNT) for synthesis or degradation. NTC pseudogenes are drawn independently so are not comparable and epistasis is not calculated.

### RPL26-3xFLAG Validation and Pulse-Chase Assay

Endogenously tagged RPL29-HaloTag screening clone containing Zim3-dCas9 was infected with RPL26-3xFLAG variants and selected with blasticidin for 20 days, freezing stocks. To confirm lack of UFMylation on 5K5R mutant, confluent cells grown at 10% serum were treated with anisomycin at 50 nM (Sigma A9789) for 15 minutes before lysis (or treated with ethanol as vehicle control), then lysed with RIPA and lysates run on 12% gel, transferred to membrane, and blotted for RPL26.

For pulldown pulse-chase assay, two replicates of cells without RPL26-3xFLAG and 3 replicates each of RPL26-3xFLAG and RPL26-3xFLAG (5K5R) were used. These cells were grown to confluence in a 10 cm dish and serum reduced to 1% for 5 days. JF549 was added to the cells at 100 nM for 1 hour, followed by washing with medium and counter-staining with JF646 (100 nM). Media was changed on day 3 of the experiment, and on day 5 cells were re-stained with JF646, then washed in ice-cold PBS and lysed in 400 µL polysome lysis buffer 1% Triton X-100 (Thermo Fisher Scientific 85111), 20 mM tris(hydroxymethyl)aminomethane (Tris) at pH 7.5 (Thermo Fisher Scientific 15567027), 5 mM magnesium chloride, 150 mM sodium chloride, 1 mM dithiothreitol, 24 U/mL Turbo DNAse (Thermo Fisher Scientific AM2238), 1x Roche cOmplete EDTA-free protease inhibitor (Millipore Sigma 4693132001), 20 U/mL Superase-In RNase Inhibitor (Thermo Fisher Scientific AM2696)). Cycloheximide was not included to allow subunit dissociation. Aliquots of lysate were preserved for immunoblot analysis, and the remainder pelleted through a 1 mL sucrose cushion in polysome buffer (1 M sucrose (Millipore Sigma S0389-500G), 20 mM Tris pH 7.5, 5 mM magnesium chloride, 150 mM sodium chloride) using a TLA110 rotor (Beckman Coulter 366735) in a Optima MAX-TL Ultracentrifuge (Beckman Coulter A95761). Sucrose was removed and the pellet resuspended in 200 µL of polysome buffer (20 mM Tris pH 7.5, 5 mM magnesium chloride, 150 mM sodium chloride).

Sample RNA was measured by qubit, aliquots saved for blotting and analysis, and 25 µg RNA was brought to 350 µL with polysome buffer. Triton X-100 was added to 0.05%. M2 FLAG magnetic beads (Millipore Sigma M8823) were pre-equilibrated in polysome buffer with 0.01% Triton X-100 (low-salt wash buffer) 3x to remove storage buffer, then 84 µL of 1:1 bead:buffer slurry was added to 25 µg ribosomes, and allowed to rotate in low-bind microcentrifuge tubes overnight. The next day, beads were bound to a magnetic rack (Sergi Lab Supplies 1005a) at 4°C, supernatant saved as flowthrough sample, and beads washed 1x in 400 µL (low-salt wash buffer, 3x in 400 µL high-salt wash buffer (same as polysome buffer but with 0.5 M sodium chloride and 0.1% Triton X-100), followed by resuspension in 400 µL low-salt wash buffer. All washes kept at 4°C, with 5 minutes rotation before bead binding to rack. 3xFLAG peptide (Millipore Sigma F4799-4MG) was diluted to 100 µg/mL in low-salt wash buffer; beads were pelleted then resuspended in 200 µL FLAG-peptide-containing solution and allowed to rotate for 1 hour to elute. Beads pelleted and samples collected.

RNA content measured to normalize samples via qubit, samples normalized in polysome buffer then run on 4–12% polyacrylamide SDS-PAGE. In-gel fluorescence used to measure JF549 and JF646 on Typhoon scanner, then proteins transferred to nitrocellulose membrane using transblot turbo and RPL26 blotted for to confirm purity.

To test for changes in turnover rate, integrated intensity in bands from relevant lanes on pulldown wells were calculated using ImageStudio (LICOR biosciences, v. 5.5.4) with local background. Log_2_ of ratio of new (JF646) and old (JF549) signal was calculated per-sample, and these were compared using Welch’s t-test within-sample.

### SILAC pulse-chase cell culture

RPE1 cell line with RPS3-HaloTag and either sgNT or sgDDRGK1 already selected with puromycin were grown for 3 passages (approximately 2 weeks) in DMEM/F12 for SILAC (Thermo Fisher Scientific 88370) with heavy lysine (^13^C_6_ ^15^N_2_, Thermo Fisher Scientific 88209) and arginine (^13^C_6_ ^15^N_4_, Thermo Fisher Scientific 89990) in 10% dialyzed FBS (Thermo Fisher Scientific 26400044) with penicillin and streptomycin (10 U/mL, Thermo Fisher Scientific 15140122). After the third passage, cells were grown in a 6 well plate to full confluence under these medium conditions, serum levels were reduced to 1% for 5 days, followed by overnight treatment with 20 µM STLC (Millipore Sigma 164739). Media was shifted to light SILAC medium with 1% dialyzed FBS (identical media but with non-heavy isotope lysine (Thermo Fisher Scientific 89987) and arginine (Thermo Fisher Scientific 89989), containing 20 µM SLTC). Samples were washed with PBS and collected by scraping at 3 and 8 days after medium shift-all samples were collected in triplicate and flash frozen with liquid nitrogen.

### Sample collection for non-SILAC proteomics

RPE1 cells (either RPS3-HaloTag clonal screening cell line or RPL29-HaloTag clonal screening cell line as described above) pre-infected and selected with puromycin for sgNT or sgDDRGK1 expression. Cells collected by scraping in PBS and flash-freezing either after reaching quiescence or during normal growth conditions at 70% confluence with 10% serum.

### Mass Spectrometry Sample Preparation

Cells were lysed in urea denaturing buffer (8M Urea, 150mM NaCl, 50mM EPPS pH8.0, containing mammalian HALT protease inhibitor cocktail (Sigma), and Phos-STOP (Sigma). Cell lysates were sonicated on ice for 10 seconds at level 5, and resultant extracts were clarified by centrifugation for 10 minutes at 15,000 rcf at 4°C. Lysates were quantified by BCA and ∼50 μg of protein was reduced with TCEP (10mM final concentration for 30 min) and alkylated with Chloroacetamide (20mM final concentration) for 30 minutes. Proteins were chloroform-methanol precipitated using the protocol in SL-TMT protocol,^170^ reconstituted in 200 mM EPPS (pH 8.5), digested by Lys-C for 2h at 37°C (1:200 w:w LysC:protein) and then by trypsin overnight at 37°C (1:100 w:w trypsin:protein). ∼25μg of protein was labeled with 62.5 µg of TMTpro for 120 min at room temperature. After labeling efficiency check, samples were quenched with hydroxylamine solution at ∼0.3% final (w/v in water), pooled, and desalted C18 solid-phase extraction (SPE) (SepPak, Waters). Pooled samples were offline fractionated with basic reverse phase liquid chromatography (bRP-LC) into a 96-well plate and combined for a total of 24 fractions^171^ before desalting using a C18 StageTip (packed with Empore C18; 3M Corporation), and subsequent LC–MS/MS analysis.

### Liquid chromatography and mass spectrometry data acquisition

Mass spectrometry data were collected using a Orbitrap Lumos or Eclipse mass spectrometer (Thermo Fisher Scientific, San Jose, CA) coupled with Neo Vanquish liquid chromatograph. Peptides were separated on a 100 μm inner diameter microcapillary column packed with ∼35 cm of Accucore C18 resin (2.6 μm, 150 Å, Thermo Fisher Scientific). For each analysis, we loaded ∼2 μg onto the column. Peptides were separated using a 90 min gradient of 5 to 29% acetonitrile in 0.125% formic acid with a flow rate of 300 nL/min. The scan sequence began with an Orbitrap MS1 spectrum with the following parameters: resolution 60K, scan range 350-1350, automatic gain control (AGC) target 100%, maximum injection time “auto,” and centroid spectrum data type. We use a cycle time of 1s for MS2 analysis which consisted of HCD high-energy collision dissociation with the following parameters: resolution 50K, AGC 200%, maximum injection time 150ms, isolation window 0.6 Th, normalized collision energy (NCE) 36%, and centroid spectrum data type. Dynamic exclusion was set to automatic. The FAIMS compensation voltages (CV) were-30,-50, and-70V.

### TMT Data Analysis

Mass spectra were converted to mzXML and monoisotopic peaks were reassigned with Monocole and then database searched using a Comet-based HT using Proteome Discoverer (v2.3.0.420 – Thermo Fisher Scientific).^172–174^ Database searching included all canonical entries from the Human reference proteome database (UniProt Swiss-Prot – 2019-01; https://ftp.uniprot.org/pub/databases/uniprot/previous_major_releases/release-2019_01/) and sequences of common contaminant proteins. Searches were performed using a 20 ppm precursor ion tolerance, and a 0.02 Da product ion tolerance for ion trap MS/MS were used. TMT tags on lysine residues and peptide N termini (304.207 Da for TMTpro) and carbamidomethylation of cysteine residues (+57.021 Da) were set as static modifications, while oxidation of methionine residues (+15.995 Da) was set as a variable modification. For SILAC experiments, heavy isotope-labeled lysine (Lys8, +8.014 Da) and arginine (Arg10, +10.008 Da) were included as variable modifications. PSMs were filtered to a 2% false discovery rate (FDR) using linear discriminant analysis using the Picked FDR method, proteins were filtered to the target 2% FDR level.^173^ For reporter ion quantification, a 0.003 Da window around the theoretical m/z of each reporter ion was scanned, and the most intense m/z was used. Peptides were filtered to include only those peptides with >200 summed signal-to-noise ratio across all TMT channels. An isolation purity of at least 0.5 (50%) in the MS1 isolation window was used for samples analyzed without online real-time searching. For each protein, the filtered peptide-spectral match TMTpro raw intensities were summed and log2 normalized to create protein quantification values (weighted average). Using protein TMT quantifications, TMT channels were normalized to the summed TMT intensities for each TMT channel.

### SILAC Degradation/Synthesis Data Analysis

Both heavy and light peptide channels were normalized by sum total of heavy and light signal for each individual sample after removal of contaminants and decoys. Average and standard error of the mean (SEM) for each condition (day and genotype) was calculated, and ratio between day 8 and day 3 calculated with SEM propagated as the root sum square of ratio of SEM to mean. Data were log_2_ transformed and difference, as well as Welch’s t-test based on combined standard errors, calculated in log space. Subcellular locations extracted based upon annotations from large-scale organelle IP data using graph-based localization-nucleolus category was folded into nucleus, and translation, stress granule, and p-body categories were added to cytosol.^175^ Categories not shown were removed. Ribosomal proteins manually annotated with reference to the human proteostasis network annotation.^168,169,176^

Statistical testing for categories in violin plots in Figure 3B was based upon Mann-Whitney U test between total proteome excluding category and category itself, with random subsampling to a maximum sample size of 50 (or category size, if smaller) to prevent p-value inflation for large categories-reported p-value are the median from 1000 random draws, Benjamini-Hochberg corrected for number of tested categories. The subset of localizations tested was chosen manually. For condition-comparison panels Supplementary Figure 4D–E, p-values represent significance of difference between genotypes in log_2_ fold change as shown in Figure 3B, not of individual per-genotype changes between days. Annotations available in Supplementary Table 8.

### Total Protein Data Analysis

Data normalized as median log_2_-transformed signal between samples. Log_2_ fold change calculated based on difference of mean log_2_ signal between samples, with p-values calculated as Welch’s t-test based on SEM of each group, with Benjamini-Hochberg false discovery rate correction. Categories compared to the rest of the proteome as described above for SILAC data using Mann Whitney U test subsampled to a maximum group size of 50. For membrane topology-related figures, UniProt identifiers were matched to UniProt release 2026_01,^177^ and Signal peptide, transmembrane, and terminus orientation were extracted from annotated features. Mitochondrial proteins were extracted from inclusion in MitoCarta 3.0; membrane proteins not annotated as mitochondrial were assumed to be secretory.^178^ UPR genes were manually annotated from literature.^93^ Final category annotations are available in Supplementary Table S9.

### Sucrose gradient / polysome profiling

For sucrose gradient samples, cells were grown at 10% FBS to full confluence and maintained at full confluence for 5 days. Fresh media was added 2 hours before sample collection to ensure robust translation. Cells were collected by washing with ice-cold PBS containing 100 µg/mL cycloheximide (CHX, Millipore Sigma C4859), followed by lysis on ice by scraping with polysome lysis buffer (1% Triton X-100 (Thermo Fisher Scientific 85111), 20 mM tris(hydroxymethyl)aminomethane (Tris) at pH 7.5 (Thermo Fisher Scientific 15567027), 5 mM magnesium chloride, 150 mM sodium chloride, 1 mM dithiothreitol, 24 U/mL Turbo DNAse (Thermo Fisher Scientific AM2238), 1x Roche cOmplete EDTA-free protease inhibitor (Millipore Sigma 4693132001), and 100 µg/mL CHX). Samples were clarified at 1600 rcf and 4°C, then flash frozen. Upon thawing, A_260_ of clarified lysate was measured via qubit and equal concentration of lysates, normalized to load 92 µg in 1 mL, loaded onto a continuous 10%–50% sucrose gradient (w/v in 20 mM Tris, 5 mM magnesium chloride, 150 mM sodium chloride, 1 mM dithiothreitol) produced using a BioComp Gradient master using in Polyclear centrifuge tubes (11 x 89 mm; Seton 7030). Polysomes were separated using a Beckman Coulter Optima XPN-80 Ultracentrifuge with an SW41 rotor at 36000 rpm for 2 hours at 4°C. Tubes were kept at 4°C until analysis, at which point they were fractionated and monitored using a Biocomp Gradient Fractionator equipped with a Triax flow cell. Data re-normalized based on loaded A_260_, which was found to better correspond to total area than qubit measurements. Data presented is representative of multiple replicates.

### SD40-UBA5 degron cell line creation and validation

RPE1 polyclonal cell lines with PiggyBac-encoded Zim3-dCas9 and AviTag-Halo-RPS24 or RPL29-Halo-AviTag were infected with lentivirus encoding EF1a SNAP-Sec61β-2A-SD40-UBA5 with sgUBA5 expressed from a U6 promoter. After 48 hours, these cells were stained for 1 hour with 1 µM cpSNAP-JF552 dye, washed with media and incubated for 1 hour without dye, and then sorted for SNAP-positivity.

To test the degree of UFMylation, cells were grown to confluence in 10% FBS and allowed to equilibrate to contact inhibition for 5 days with regular media changes. PT-179 (20 µM from 40 mM stock solution in DMSO, Selleck E1937) or equivalent amounts of DMSO were added for indicated amount of time. To induce UFMylation, 50 nM anisomycin was added for 10 minutes, followed by lysis in RIPA buffer and immunoblot on nitrocellulose for UFM1 and GAPDH, imaged on LICOR imager and quantified using ImageStudio with local background.

Quantification is of specific band regions seen for mono-and di-UFMylation of RPL26, and normalized to GAPDH signal.

Flow-chase assays were performed as described in section “HaloTag pulse-chase turnover assay by flow cytometry” with pre-treatment with PT-179 (20 µM) for indicated amounts of time.

### Single-molecule tracking data acquisition

The majority of single-particle tracking experiments used Avi-Halo-RPS24 and RPL29-Halo-Avi constructs as described above, integrated via lentiviral infection into polyclonal RPE1 cells with CAG Zim3-dCas9 and FACS sorted for cells positive after staining with JF646 HaloTag dye. Experiments in Supplementary Figures 6D–E used endogenously editing RPL29-HaloTag screening cell lines with sgNT or sgUBA5 guide RNAs. Cells were grown to confluence in 24-well glass coverslip plates (Cellvis P24-0-N) at 10% serum for all experiments. To achieve sub-stochiometric labeling of ribosomes (to enable distinguishing individual particles), Janelia Flour X 650-HaloTag ligand (JFX650) was prepared at 100 nM, and 10 µM 1-chloro-6-(2-propoxyethoxy)hexane (CPXH) was prepared as a nonfluorescent blocker,^179^ both in DMEM/F12 at 10% FBS. JFX650 solution was then diluted 1:1000 in CPXH solution, and this mixture was added to cells in multiwell plate, incubated for 1 hour, then media was removed, rinsed 1x with fresh medium, and allowed to recover in fresh medium for at least 1 hour but no more than 7 hours prior to imaging. For ER-coimaging experiment, Janelia Fluor 552-cpSNAP-tag was added at 1 µM to degron cells described above at time of ribosome labeling; UBA5 degradation was induced 24 hours prior to imaging with addition of 20 µM PT179, or equivalent amounts of DMSO (0.1%) in control cells, and drug concentration was maintained during and after ribosome and SNAP-tag labeling. For polysome run-off/stabilization experiments, harringtonine (MedChemExpress, HY-N0862) was added at 10 µg/mL in complete media for 1 hour prior to imaging; emetine dihydrochloride (Millipore Sigma, E2375) was added to 360 µM in complete media for 1 hour prior to imaging.

Single-molecule tracking experiments were conducted as previously described^180^ on a custom-built Nikon (Nikon Instruments Inc.) Ti-2E microscope equipped with a 100x (NA 1.49) oil-immersion TIRF objective (Nikon apochromat CFI SR HP Apo TIRF 100x Oil), two Prime 95B sCMOS cameras (Teledyne Photometrics), a perfect focus system to correct for axial drift and motorized dual galvo laser illumination system (iLas2, Gataca Systems), which allows an incident angle adjustment to achieve highly inclined and laminated optical sheet illumination.

The incubation chamber maintained a humidified 37 °C atmosphere with 5% CO2 and the objective was similarly heated to 37 °C for live-cell experiments. 1 hour drug additions were performed on microscope and allowed to equilibrate. Excitation was achieved using a 640 nm laser line (1 W, Coherent Genesis) for JFX650, and with a 561 nm laser line (1 W, Coherent Genesis) for JF552 excitation. The excitation lasers were modulated by an acousto-optic Tunable Filter (AOTFnC-400.650-TN, AA Opto-Electronic). The laser light is coupled into the microscope by an optical fiber and then reflected using a multiband dichroic (405 nm/488 nm/561 nm/633 nm quad-band, Semrock) and then focused in the back focal plane of the objective. Fluorescence emission light was split using a 635 nm dichroic mirror, then filtered using either a 593/40 nm or a 698/70 nm band-pass filter placed in front of the camera. The microscope, cameras, and hardware were controlled through the NIS-Elements software (Nikon).

All single-particle tracking data were acquired under constant illumination unless otherwise noted below. Data presented in Figure 5 and Supplementary Figure 6 was collected with 15 ms frame interval, pixel size 110 nm. For Figure 5C and Supplementary Figures 6A–C, 1000 frame imaging was performed on 10 fields of view per condition (1024×704 pixels, 110 nm pixel size). For Supplementary Figures 6D–E, 500 frames of imaging were performed on 20 fields of view per condition (532×348 pixels, 110 nm pixel size). For ER-imaging experiments, 100×15 ms frames of constant 561 nm illumination were captured with sCMOS with 593/40 nm bandpass, followed by 1000 frames of constant 640 nm illumination captured on, for 10 fields of view for the DMSO condition and 14 for the PT-179 treated condition (800×700 pixels, 110 nm pixel size).

### Single-molecule tracking data analysis

All analysis in python 3.8. Spots were detected and localized with Quot (commit 1b9051e, Jan 25 2024, <u>github.com/alecheckert/quot</u>); particles were detected by log-likelihood-ratio detection followed by least-square fitting of a 2D gaussian, and localizations were linked by Euclidean nearest-neighbors with a 1.2 µm maximum radius and no gap closing. Diffusion distribution was estimated using saSPT v 0.4.0 in regular-Brownian-motion-with error (RBME) mode, marginalizing over a state array of 150 diffusion states from 10^-3^–10^2^ evenly spaced in log_10_ crossed with 36-state localization error states from 0–0.070 in 0.002 µm steps (the saSPT default) with trajectories split to at most 10 displacements, with 200 variational Bayes expectation maximization iterations. Focal depth was set to 0.7 µm throughout; conclusions were insensitive to focal depth. Trajectories for all fields of view in a condition were combined before inference to maximize statistical power. Full saSPT and quot parameters are listed in Supplementary Table 12.

Diffusion bins were chosen by manual inspection of data across datasets, to represent consistent troughs. Bins likely contain multiple molecular species that cannot be fully resolved via the current analysis.

“Probability density” in figure legends is posterior occupancy after saSPT inference, and represents the normalized weighted sum of posterior probability distributions from each measured trajectory; it represents a probability density per unit in log_10_ space. “ΔProbability Fraction” is the change in integrated density within a diffusion bin-since bins are not the same length, total change magnitude cannot be accurately compared across bins.

Due to Bayesian nature of analysis potentially changing estimated diffusion coefficient of any single trajectory if combined with different fields of view, all statistical analysis was done via re-iteration of saSPT with altered sets of fields of view; 500 iterations were used for all resampling.

ΔProbability fraction represents differences in integrated density in a bin. All 95% confidence intervals were established via bootstrap resampling of both conditions with replacement of fields of view within a condition, followed by measurement of difference in probability density in a given bin. estimated based on the central 95% (475) resamples. p-values for contrasts between conditions were established via random permutation of fields of view-p-values represent (1 + k)/(501), ie number of null draws more extreme than initial metric plus one divided by total draws plus one, setting a floor for p-value of 0.001996.

### ER tracking analysis

Median projection of 100 frames of ER tracking were illumination-corrected by dividing by an image blurred with a 100 pixel Gaussian, multiplied by original image mean; this is the example image shown in Figure 5E. Illumination correction was repeated identically to improve contrast, and images were subjected to two iterations of Richardson-Lucy deconvolution with a 0.74 pixel gaussian point spread function, renormalized to mean, before thresholding for mask construction.

ER mask was constructed in two stages, using local Gaussian-weighted block mean. Mask inclusion could either pass a local threshold of 1.5% above Gaussian weighted mean with 25 pixel radius or the same 1.5% local thresholding but with a 101 pixel radius. Both separate radii masks were subjected to disk closing with a radius of two pixels to remove noise and link nearby objects, contiguous objects smaller than 500 pixels were removed, and any pixels less than the 60% for an image were removed. Both masks were then combined for the final ER mask.

To establish a trajectory as “on-ER”, each trajectory was tested for enrichment on this mask using a binomial enrichment test with a threshold of p<0.05 for categorization. To accommodate the large off-ER area, trajectories were only considered off-ER if they are >95% of localizations outside the ER mask and had a minimum number of localizations required to pass the binomial test (which depends on per-FOV mask percent area). Shorter trajectories and those with mixed localization are considered “undetermined” and not analyzed in the ER analysis. All labeling was performed at the trajectory level before choosing subtrajectories with saSPT; saSPT was independently fit as described above to derive the diffusion distribution. Note: we do not represent this masking procedure as ground truth for the ER, but as a confident enrichment for ER trajectories to allow comparison.

saSPT and resampling for Figure 5F was performed only on the trajectories passing this mask. For confidence intervals, field of view bootstrap resampling was performed as described above. From comparison between treatments, all data were jointly fit in a single saSPT analysis then re-split by treatment and the difference-in-differences of “on ER” – “off ER” averaged for PT-179 FOVs minus the same averaged difference for DMSO treated FOVs. The FOVs are then randomly permuted to construct a null distribution, repeating this procedure; because only one saSPT run is performed, 10,000 permutations are able to be done, setting a lower p-value floor for this comparison.

### Optogenetic proximity labeling (LOV-BirA) and chase analysis

RPE1 polyclonal cell lines with PiggyBac-encoded Zim3-dCas9 and AviTag-Halo-RPS24 or RPL29-Halo-AviTag were infected with lentivirus encoding pSFFV SNAPtag-2A-V5-LOV-BirA-Sec61β. After 48 hours, these cells were stained for 1 hour with 1 µM cpSNAP-JF552 dye, washed with media and incubated for 1 hour without dye, and then sorted for SNAP-positivity. Sorted cell populations infected with guide RNAs against relevant genes, selected, then grown in dark room without exposure to non-red light for at least two passages to remove any residual background biotinylation; one extra plate was plated for monitoring confluence in non-light conditions. Pulse-chase assays carried out in two formats as described below:

### Flow cytometry

Cells plated in 24-well plates with 12 wells per genotype plated for each timepoint and light condition (immediate light, immediate no light, 4 days no light) plated on a separate plate, combining genotypes. All operations were carried out in a dark room under red light illumination through permeabilization. After reaching confluence, cells were maintained for 5 days. All well numbers below are per-genotype. For 4-day chase samples, 9 wells were labeled with JF549 HaloTag dye, and 3 wells each initially left unstained. After staining, wells were washed, and media with 5 µM supplemental biotin (Millipore Sigma B4639-100MG) was added. Plate was placed on a blue light box (Major Science MBE-200BW) inside a humidified incubator at 5% CO_2_ and 37°C and exposed to light for 18 minutes. After light exposure, media was removed and media without supplemental biotin containing to JF646 (100 nM HaloTag dye) was added to the 3 previously-unstained wells and 6 previously-JF549-stained wells; three of the previously-JF549-stained wells received media without dye as singly stained controls. Cells were returned to incubator, and media changed after 24 hours to dye-free media. After 4 days, day 4 chase samples were re-stained with their final color, giving 6 chase wells and 3 singly-stained controls per dye; both light and dark day 0 samples had 4 wells each staining with JF646, JF549, or lack of staining as control. After staining for 1 hour, dye was removed, washed with media, and 5 µM biotin was added to day 0 samples. Day 0 “light” samples were exposed to light for 18 minutes. At end of this, all plates were washed with PBS, dissociated with TrypLE, triturated to finish dissociation, TrypLE quenched with media, and samples transferred to 1.5 mL centrifuge tubes. Cells were pelleted at 500 rcf at 4°C, washed 400 µL PBS, and fixed for 1 hour at 4°C with 200 µL 4% paraformaldehyde in PBS. After fixation, 1 mL of FACS buffer (1% BSA in PBS with 1 mM EDTA) was added to each tube, triturated, and cells pelleted at 1000 rcf. Cells were then permeabilized with 0.1% Triton X-100 in FACS buffer for 20 minutes, followed by washing with FACS buffer. Cells were resuspended in 100 µL FACS buffer with 2 µg/mL streptavidin-AlexaFluor 568 (reconstituted in PBS, Thermo Fisher Scientific S11226) and incubated for 1 hour, after which cells were pelleted, buffer removed, washed 3x with 5 minutes incubations at room temperature to remove excess streptavidin. During labeling, fixation, and staining, singly-stained controls were prepared for compensation. Final samples were resuspend in 200 µL FACS buffer and transferred to 96 well plate.

Samples were acquired using attune flow cytometer with autosampler, with singly-stained controls used to set up compensation settings according to manufacturer’s protocols. Median compensated fluorescence (MFI) after gating for single-cells, GFP (guide vector), and BFP positivity was calculated, and all samples were normalized per-genotype by subtracting the average MFI for all unstained samples (or, for biotinylation, not-exposed-to light, to account for endogenous biotinylation background), followed by dividing by the average MFI for non-chase fully-stained samples to normalize per-genotype between 1 (fully labeled) and 0 (unlabeled).

Samples with multiple staining patterns were combined for totals, giving variable numbers due to different staining patterns between immediate labeling and chase samples. For JF549 samples, we noted light exposure likely bleached a proportion of signal (70% of dark samples)-to account for this dark samples were not included in calculation of fully-labeled averages. Statistical comparison of normalized values was based on Welch’s t-test. Only biotinylated samples passed a p<0.05 level of significance.

### In-gel fluorescence

Cells maintained as described above, but all samples maintained in 6-well plates-each plate contained all 6 relevant genetic manipulations, and 6 replicates were plated in triplicate for dark labeling, light labeling, and 5-day chase samples. Upon reaching confluence, media was shifted to 1% FBS for 5 days prior to labeling any samples. At time of labeling, 500 nM biotin was added to all samples, and all except dark samples were co-illuminated on light box (465 nm wavelength, 14-Watt Power blue light, Amazon, model number 884667106091218, discontinued) for 15 minutes in humidified incubator. We note day 0 and day 5 samples were illuminated at the same time. Media was removed, and all samples were stained with JF549 100 nM (HaloTag dye) for 1 hour. After labeling, 5-day chase samples were washed with dye-free media and counter-stained with CPXH 10 µM, left for 24 hours before removal and replacement with dye-free media. For immediate timepoints with and without light exposure, after 1 hour with JF549 these samples were washed with PBS and lysed with 200 µL RIPA as described for immunoblot preparation in dark room on ice; samples were clarified and supernatant was added to 50 µL 10% SDS to fully denature, then flash frozen. For 5-day chase samples, cells were lysed after 5 days of incubation as described. After lysis, all samples were run on 4–12% Bolt bis-tris gels by SDS-PAGE as described under section “SDS-PAGE and Immunoblot Protocols”. 16 samples were loaded per gel (along with a ladder), with single replicates of, leading to replicates spread across 8 gels along with a ladder; we note one replicate of sgNT and sgUBA5 was run twice due to extra lanes, but excluded from analysis. In gel fluorescence was collected via Typhoon scanner, then proteins transferred to PVDF membranes via high MW program on transblot turbo, stained for HaloTag and streptavidin (700 nm wavelength) overnight, stained with secondary antibody (800 nm) against HaloTag for 1 hour, then imaged as a single set of membranes at fixed scanning conditions, with washing as described. Membranes were then re-probed with anti-tubulin antibody and secondary antibody (800 nm), and re-imaged as a single set.

For data analysis, all lanes were analyzed using ImageStudio with local background from above and below band. Tubulin-probed rescan is only used for tubulin quantification. To account for any differences in loading, all band intensities normalized to tubulin, and day 0 sgNT light-exposed samples for biotin staining, or mean of dark and light exposed samples from day 0 for sgNT for JF549 levels. For autophagy level quantification, we quantified the ∼30 kDa lower band that has been previously reported to be autophagy dependent; this band is only present in the 5 day chase samples, consistent with stabilization requiring binding to rhodamine-based dyes (like the JaneliaFluor dyes). For space and the need to adjust contrast to see the upper intact band and lower non-intact band. No biotin signal was seen on these lower bands, consistent with only the stable core of HaloTag surviving in the lysosome; the reduced turnover rate upon loss of lysosome activity suggests this extended lifetime is limited. We note the presence of an intermediate ∼40 kDa band that we take to be post-lysis proteolysis of the HaloTag, that highly correlates with intact band intensity for samples lysed on the same day, but with reduced levels for the 5 day samples based on HaloTag quantification. We included both intact and this intermediate cleavage bands in all quantifications of intact HaloTag; these make the differences in HaloTag-per-tubulin values negligible between collection types.Total ribosome quantification based on JF549 (for day 0 samples) or HaloTag gave similar results, so JF549 was analyzed for consistency. Displayed images represent a single replicate from a single plate; discontinuity is due to samples being on separate gels, stained alongside samples from a different replicate, confirming effects are not due to membrane-to-mebrane staining variability. All intensities analyzed via Welch’s t-test with Benjamini-Hochberg correction for multiple testing (the subset of samples compared are indicated in figure legends). 95% confidence intervals were calculated based upon the t-distribution with 5 degrees of freedom.

### RPL29-mKeima ribophagy assay

RPL29-mKeima was infected into both polyclonal RPE1 cells with PiggyBac Zim3-dCas9 as well as these same cells already infected and sorted for expression of degradable EF1a SNAP-Sec61β-2A-SD40-UBA5 construct. Since these cells have reduced baseline UFMylation before degradation (Supplementary Figure S6), ribophagy levels were measured in both the degron and non-degron cells. All samples were grown to confluence in 24-well plates with n=6 samples each for Bafilomycin A1 treated vs untreated and EBSS-starved vs. unstarved, giving 24 plated samples per treatment. PT-179-treated samples were treated with 20 µM PT-179 for 6 days prior to starvation, 7 days at time of collection, and new media with or without compound was added every two to three days. 20 hours before collection, 12 wells of each sample were shifted to amino-acid-free EBSS, and half of both starved and non-starved samples (6 wells per condition) were treated with Bafilomycin A1 at 10 nM. Samples collected at 24 hours via dissociation in TrypLE and running on attune, with VL3 (excitation 405 nm, emission 603/48 nm bandpass) as neutral detection and YL2 (excitation 561 nm, emission 620/15 nm bandpass) used to collect acidic signal. Median fluorescence intensity after gating on BFP positivity and single cells was calculated for each. Due to prolonged Bafilomycin treatment, increases in neutral species, which we presumed to be due to feedback inhibition of proteasomal degradation of unincorporated ribosomal proteins. We therefore measured change upon Bafilomycin based upon median fluorescence intensity, with standard error for treated and untreated combined; data shown in Figure 6C and Supplementary Figure 7D for acidic and neutral, with Welch’s t-test (Benjamini-Hochberg corrected p-values) used to compare between samples.

### Confocal colocalization of GFP-TLSD1 with the ER

GFP-3xHA-TLSD1 lentivirus was used to infect RPL29-HaloTag-Avi cells containing the sgUBA5 construct expressing SD40-UBA5, to allow temporal control of UFMylation. Cell population was sorted for GFP and BFP positivity to ensure continued expression of PiggyBac encoded Zim3-dCas9 machinery. This polyclonal cell line was plated on 24-well glass coverslip plates (Cellvis P24-0-N) and grown to moderate confluence with 10% serum. PT179 (20 µM) was added 13 hours before imaging from 40 mM stock; equivalent amounts of DMSO were added to control wells. Before imaging SNAP-JF552 1 µM(kind gift from Luke Lavis) and JFX650-HaloTag (100 nM) dyes was added to cells for 1 hour; cells were washed with media, then left to incubate in media for 1 hour before imaging. Imaging was performed on a customized Nikon Crest V3 spinning disk confocal microscope. (Nikon Instruments Inc.). In brief, this system includes a Ti2-E automated microscope with 40x water objective (CFI APO LWD 40X WI LAMBDA S, NA=1.15) with a pixel size of; an automated stage for x-y axes movement and a piezo for z-axis movement (Queensgate); a Orca-Fusion BT sCmos camera (Hamamatsu); and a 7-channel-laser illumination system with digital control and compatible excitation/emission filters, which yields up to 1W for each channels (Lumencor Inc.). The microscope, camera, laser and other hardwares were controlled through Nikon’s NIS-Element software (Nikon). Channels were imaged sequentially with GFP excited by 488 nm laser at 100% power (500 ms exposure, FITC filter), SNAP-JF552 ER marker was excited by 545 nm laser at 50% power (1 second exposure, Cy3 Filter), and JFX650 ribosomes imaged with 640 nm laser at 50% power (400 ms exposure, Cy5 Filter), all with appropriate emission filters; perfect focus was engaged to maintain focus, and 43 z-stacks with 0.4 µm spacing were used per sample to span cell layer. To further boost the contrast, images were further enhanced by the denoise-AI function in NIS-Element software after acquisition using batch processing.

For analysis, custom python scripts (Python 3.10) used CellPose-SAM v4.09^181^ (PyTorch 2.10.0) to segment cells on GPUs on Whitehead Institute computing cluster. Because no nuclear or membrane image was present, based upon combination of channels, scaled to its own 99.9th percentile intensity and clipped to range [0,1]; per-pixel max scaled intensity between the three samples was input to make a mask, with the assumption that the combination of ribosomes, ER, and GFP would provide dense cytoplasmic and nuclear signal. Segmentation was run slice by slice in 2D (diameter = 260 px as approximated from manual measurements, flow threshold = 0.4, cell-probability threshold = 0), and slices stitched into 3D objects using Cellpose’s IoU stitching (threshold 0.5). Cellpose was used to fill holes and discard objects smaller than 500 voxels. This yielded 276 cells for DMSO condition and 192 cells for PT-179 condition. For each image, each channel was clipped between the 0.1 and 99.9 percentile of nonzero voxels to limit the influence of individual high voxels. Pearson’s correlation coefficient r was computed between each pair of channels across all voxels inside each cell’s 3D mask using scipy 1.15.2. Other packages used in data processing include nd2 0.11.2, NumPy 2.2.6, scikit-image 0.25.2, pandas 2.3.3, tifffile 2025.5.10. matplotlib 3.10.8 and seaborn 0.13.2 used for plotting. GFP-R3HDM4 and ER stain compared using Mann-Whitney U test.

### Quantification and Statistical Analysis

All relevant statistical tests are described alongside analysis in individual methods sections, with brief descriptions in figure legends.

